# A disease associated mutant reveals how Ltv1 orchestrates RP assembly and rRNA folding of the small ribosomal subunit head

**DOI:** 10.1101/2023.07.10.548325

**Authors:** Ebba K. Blomqvist, Haina Huang, Katrin Karbstein

## Abstract

Ribosomes are complex macromolecules assembled from 4 rRNAs and 79 ribosomal proteins (RPs). Their assembly is organized in a highly hierarchical manner, which is thought to avoid dead-end pathways, thereby enabling efficient assembly of ribosomes in the large quantities needed for healthy cellular growth. Moreover, hierarchical assembly also can help ensure that each RP is included in the mature ribosome. Nonetheless, how this hierarchy is achieved remains unknown, beyond the examples that depend on direct RP-RP interactions, which account for only a fraction of the observed dependencies. Using assembly of the small subunit head and a disease-associated mutation in the assembly factor Ltv1 as a model system, we dissect here how the hierarchy in RP binding is constructed. Our data demonstrate that the LIPHAK-disease-associated Ltv1 mutation leads to global defects in head assembly, which are explained by direct binding of Ltv1 to 5 out of 15 RPs, and indirect effects that affect 4 additional RPs. These indirect effects are mediated by conformational transitions in the nascent subunit that are regulated by Ltv1. Mechanistically, Ltv1 aids the recruitment of some RPs via direct protein-protein interactions, but surprisingly also delays the recruitment of other RPs. Delayed binding of key RPs also delays the acquisition of RNA structure that is stabilized by these proteins. Finally, our data also indicate direct roles for Ltv1 in chaperoning the folding of a key rRNA structural element, the three-helix junction j34-35-38. Thus, Ltv1 plays critical roles in organizing the order of both RP binding to rRNA and rRNA folding, thereby enabling efficient 40S subunit assembly.

## Introduction

In all forms of life, ribosomes are responsible not just for producing the right amount of the correct protein, but also the detection of damaged mRNAs. Both functions depend on the functional integrity of the ribosome pool, as well as efficient ribosome assembly, as reduced ribosome numbers can affect both mRNA selection [1, 2], as well as its surveillance for damage [3].

In eukaryotes, ribosomes are assembled from their constituent 4 ribosomal RNAs (rRNAs) and 79 ribosomal proteins (RPs) with the help of ∼ 200 assembly factors (AFs), which are largely conserved from yeast to humans. These factors integrate rRNA processing with binding of RPs, chaperone rRNA folding, allow for regulation, prevent premature translation initiation, and facilitate quality control [4–6].

The binding of ribosomal proteins (RPs) to ribosomal RNA (rRNA) is very hierarchical, as first characterized by Nomura and Nierhaus [7–9]. A potential advantage of this hierarchy is the ability to ensure that every RP is incorporated into the nascent ribosome, as the next step cannot proceed without the previous one. This is critical, as demonstrated by findings that individual RPs are often depleted in cancer cells, which is associated with a poor prognosis [10–13]. In addition, hierarchical binding can help avoid dead-end pathways, which limit the efficiency of assembly. While contacts between an earlier and a later-binding RP can explain some of the hierarchy (e.g., the dependence of Rps19 binding on prior binding of Rps16 and Rps18, **Figure 1A**), other parts of the hierarchy are not explained in this manner (e.g., why Rps25 is recruited *after* Rps19, **Figure 1A**). Moreover, it appears that the recruitment of some RPs, like Rps31, might be delayed, as it occurs only much after the binding if its only binding partner, Rps12 (**Figure 1A)**. Thus, the molecular forces underpinning the hierarchical assembly of RPs remain unclear.

**Figure 1:**
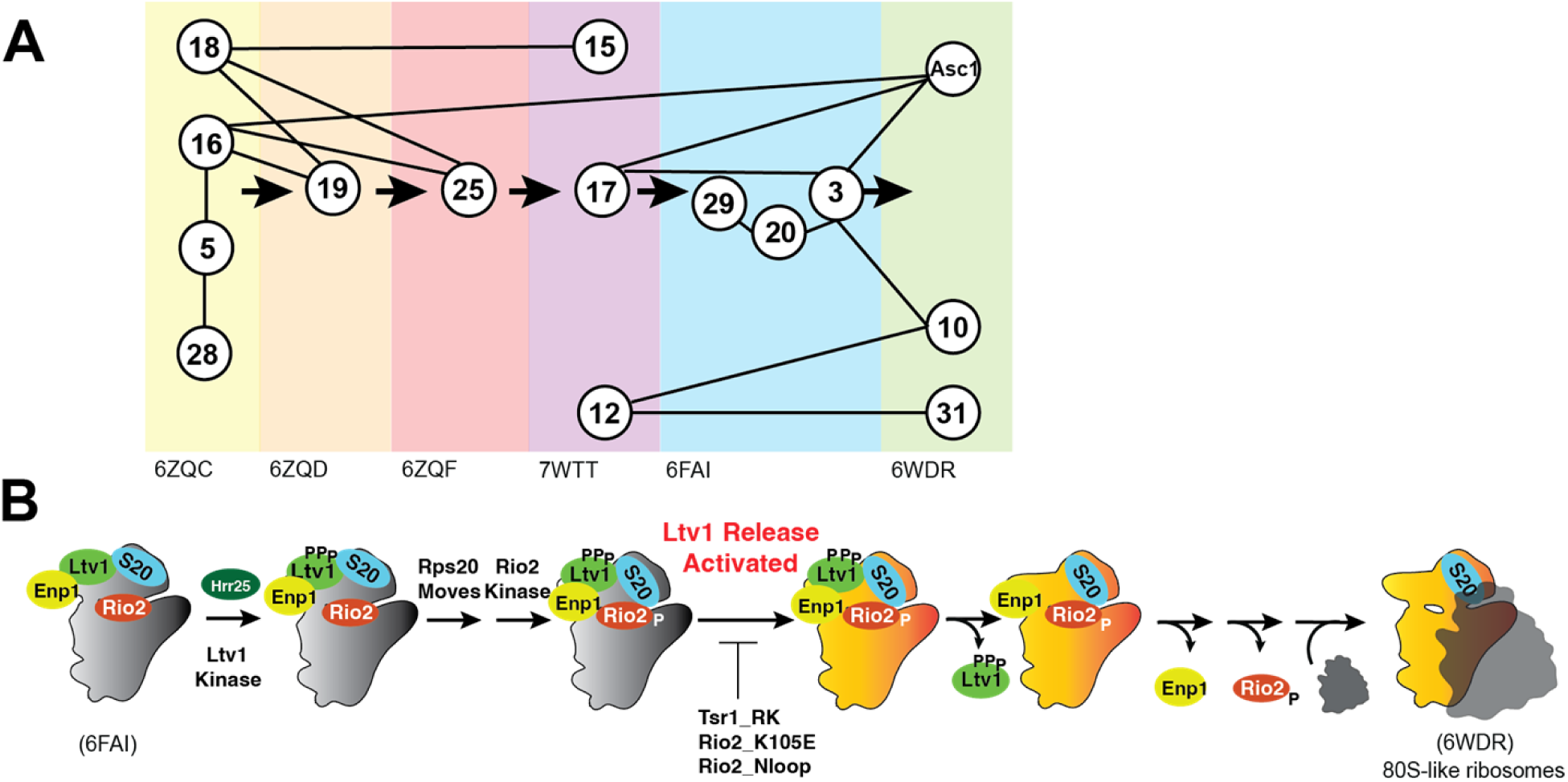
Assembly of the small subunit head is hierarchical. (A) Summary of the assembly hierarchy of ribosomal proteins to the head of the small ribosomal subunit. Assembly order was garnered from cryo-EM structures (indicated with the PDB ID at the bottom). Each intermediate is highlighted with a different color. Ribosomal proteins are indicated with the eukaryotic nomenclature. Contacts between different ribosomal proteins are indicated with line connections between the proteins, and arrows denote assembly progress. (B) A hierarchical set of steps regulates the formation of 80S-like ribosomes. Adapted from [21]. This cascade is initiated from stable pre-40S intermediate via phosphorylation of Ltv1 by the kinase Hrr25. While phosphorylation is necessary for Ltv1 release, it is not sufficient, and temporally separated. An unknown step that activates Ltv1 release was previously identified via its sensitivity to the Tsr1_RK, Rio2_K105E and Rio2_loop mutation [21].

The small subunit head is constructed from the 18S rRNA 3’-domain and is the last of the subunit substructures to emerge. Because of its relatively late assembly, individual assembly steps can be identified using cryo-electron microscopy (cryo-EM, [14–19]), as well as biochemical investigations [20–25], rendering the head a model system to study rRNA folding [22] and RP binding *in vivo* [20–24, 26]. Its assembly is initiated co-transcriptionally with binding of Rps5 (uS2), Rps16 (uS9), Rps18 (uS13) and Rps28 [27], and then continues with binding of Rps12, Rps15 (uS19), Rps17, Rps19, Rps25 [19]. The final proteins to assemble onto the head are Rps3 (uS3), Rps10, Rps20 (uS10), Rps29 (uS14), Rps31 and Asc1 [25]. While these three groups essentially recapitulate Nomura’s description of early, middle and late binding proteins [8], a more detailed analysis of the available structural evidence indicates a more intricate assembly pathway with additional hierarchies (**Figure 1A)**. Importantly, while the original dependencies could all be explained by direct binding interactions, the picture that emerges from structural work is more complex. For example, Rps19 and Rps25 both form direct interactions with Rps16 and Rps18. Yet, Rps19 binds prior to Rps25. Similarly, Rps31 recruitment is delayed until 80S-like ribosomes are formed, even though its binding partner Rps12 is present earlier. How this hierarchical assembly pathway arises remains unknown.

The assembly factor Ltv1 has emerged as a critical player in 40S head assembly [20, 21, 24, 26, 28, 29]. Binding directly to Rps3, Rps15, and Rps20 [24, 29–31], it also plays critical roles in assembly of Rps10 and Asc1 and was implicated in the folding of a critical 3-helix rRNA junction, j34-35-38 [14, 15, 22]. Moreover, it is a key actor in quality control of head assembly, which follows a hierarchical set of events that include the phosphorylation-mediated dissociation of Ltv1 (**Figure 1B**, [21]). Notably, even though Ltv1 phosphorylation is necessary for its dissociation, it is temporally separated, and first followed by rearrangements of Rps20, which in turn enable Rio2 phosphorylation. Rio2 phosphorylation then allows for an undefined step that activates Ltv1 release. Ltv1 release leads to folding of j34-35-38, dissociation of Enp1 and then Rio2, and ultimately the formation of 80S-like ribosomes, critical quality control intermediates that couple the final maturation steps to quality control. Importantly, this hierarchical sequence of events depends on the correct binding and positioning of Rps3, Rps15, and Rps20 as well as the correct folding of j34-35-38, and ensures that the newly-made subunits can accurately discriminate against non-AUG start codons [21]. Nevertheless, the full range of Ltv1 functions as well as their mechanistic basis has remained unclear as all studies have been carried out with Ltv1 deletion mutants, which tend to obscure later phenotypes.

Using a mutation in Ltv1 that causes a rare dermatological condition, LIPHAK syndrome [32], as well as a collection of genetic, biochemical, structural and mass spectrometry analyses, we demonstrate that Ltv1 plays global roles in assembly of the small subunit head, as nearly all RPs are depleted from this structure in ribosomes from cells with the LIPHAK mutation. DMS-MaP Seq was used to identify regions of change, and this analysis, combined with genetic and biochemical analyses demonstrates how Ltv1 establishes the hierarchy in RP binding, by delaying critical contacts between Rps3 and Rps20 to enable the recruitment of Rps29, and by regulating the recruitment of Rps31. Thus, our mechanistic analysis explains how incorporation of Rps29 and Rps31 is managed and explain why these are the two proteins most depleted from the subunit in cells with the disease-associated Ltv1_L216S. Thus, Ltv1 is a master regulator of assembly of the small ribosomal subunit head, with roles both in assembly and quality control.

## Results

### The disease-associated Ltv1_L216S mutation confers a growth phenotype in yeast

Recently, the same mutation in human Ltv1, Ltv1_N168S was recovered in two unrelated families suffering from LIPHAK syndrome, a rare dermatological disorder characterized by altered skin pigmentation arising from subcutaneous oxygenation defects [32]. We reasoned that this mutation could be used to uncover functions for Ltv1 in 40S ribosome assembly beyond those identified from the absence of the protein, which leads to an rRNA folding defect [21, 22, 26] that might mask additional deficiencies. We therefore set out to use our established yeast system to study defects in assembly that result from this mutation. Sequence alignments show that N168 in humans assumes the same position as L216 in yeast (**Figure S1A**). We therefore used our previously described Ltv1 deletion strain and plasmids encoding either wild type (wt) Ltv1 or Ltv1_L216S under the constitutive TEF promoter to test whether Ltv1_L216S confers a growth defect in yeast. Quantitative growth measurements show that indeed Ltv1_L216S produces a small growth defect (**Figure 2A**), which is specific for Ltv1_L216S and not observed for the neighboring Ltv1_L217S (**Figure S1B**). The magnitude of the growth defect is the same when Ltv1 is produced from the strong TEF promoter, and the weaker Cyc1 promoter, indicating that the defect does not arise from weak Ltv1 binding or low levels of the mutant Ltv1 protein (**Figure S1C**), consistent with Western blot analyses that show the same levels of wild type and mutant Ltv1 (**Figure S1D**). Thus, the disease-associated Ltv1_L216S mutation (LIPHAK mutation) causes a growth defect in yeast.

**Figure 2:**
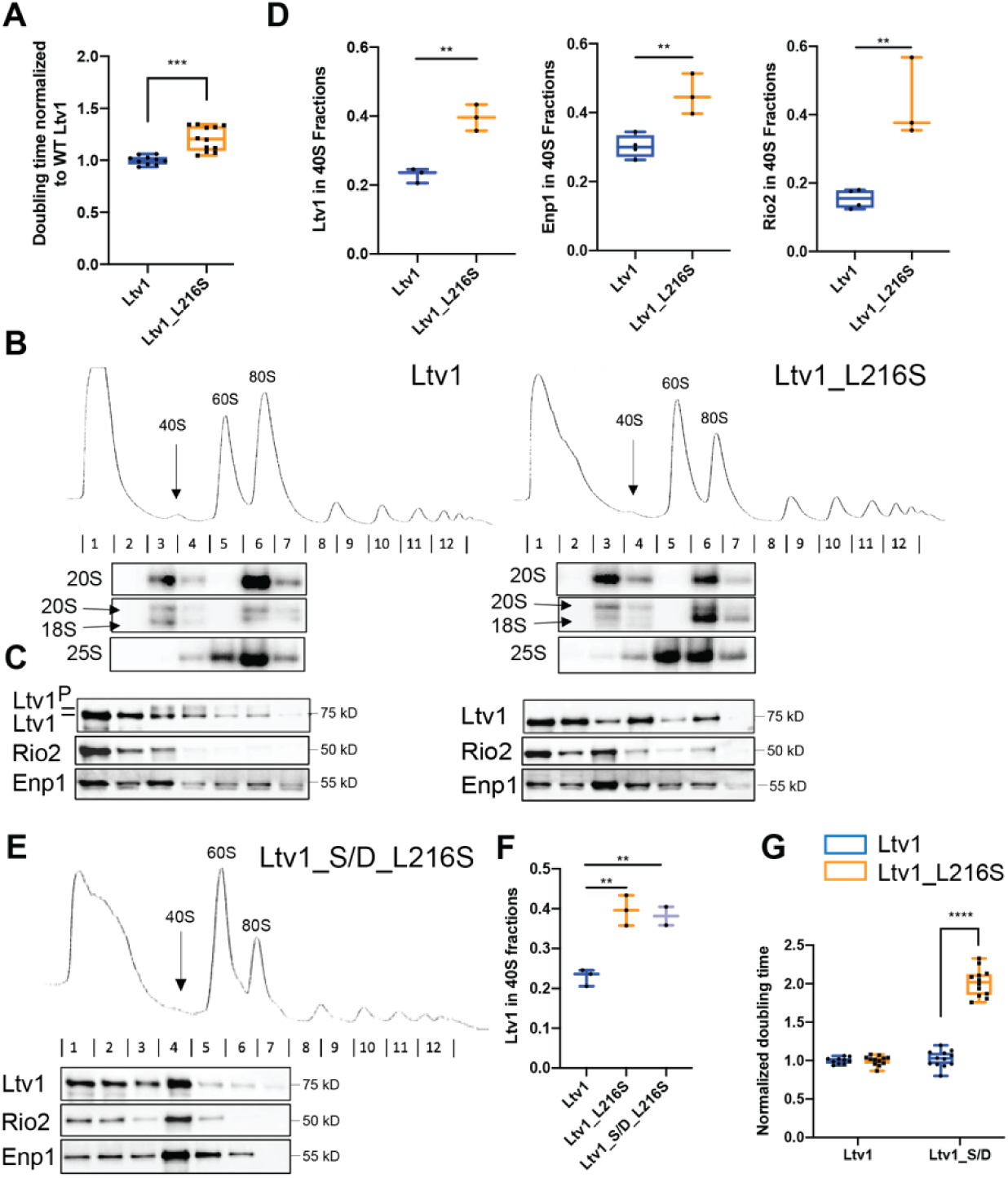
The LIPHAK-syndrome associated Ltv1_L216S mutation affects 40S head assembly, by impairing Ltv1 phosphorylation. (A) The Ltv1_L216S mutation produces a small growth defect. Growth of yeast strains lacking endogenous Ltv1 and expressing plasmid-encoded Ltv1 from the TEF promoter was measured, and doubling times extracted. Significance was tested using an unpaired t-test. ***, P<0.001. (B) Absorbance profiles at 260 nm of 10-50% sucrose gradients from Fap7 depleted cells expressing only plasmid-encoded wt Ltv1 or Ltv1_L216S. Northern blots of each gradient fraction are shown. (C) Western blots for Ltv1, Enp1 and Rio2 are shown. The position of phosphorylated Ltv1 is indicated. (D) Quantification of the data in panel C and biological and technical replicates, to demonstrate accumulation of Ltv1 (left), Enp1 (middle) and Rio2 (right) in pre-40S ribosomes. Significance was tested using an unpaired t-test. **, P<0.01; ***, P<0.001 (E) Absorbance profiles at 260 nm of 10-50% sucrose gradients from Fap7 depleted cells expressing only plasmid-encoded wt Ltv1 or Ltv1_S/D_L216S. Western blots for Ltv1, Enp1 and Rio2 are shown. (F) Quantification of the data in panel E and biological and technical replicates, to demonstrate accumulation of Ltv1 in pre-40S ribosomes. Significance was tested using an unpaired t-test. **, P<0.01. Note that data for WT Ltv1 and Ltv1_L216S are the same as in panel D and replotted for comparison. (G) Normalized (to WT Ltv1) doubling times from yeast expressing wt Ltv1, Ltv1_L216S, Ltv1_S/D or Ltv1_S/D_L216S demonstrate that the phosphomimetic Ltv1_S/D mutation [20] does not rescue, but instead exacerbates the effect from Ltv1_L216S. Significance was tested using an unpaired t-test. ****, P<0.0001.

### Ltv1_L216S is mispositioned blocking its phosphorylation-dependent release

Ltv1 is a late-binding and acting 40S ribosome assembly factor. Thus, we wanted to confirm that the growth defect from the Ltv1_L216S mutation reflected a ribosome assembly defect, as expected. Absence of Ltv1 leads to defects in rRNA folding and recruitment of late-binding proteins Rps10 and Asc1 and reduces 40S ribosome levels [21, 22, 26]. Mutations that block its release are dominant negative [20], and impair the formation of so called-80S like ribosomes. 80S-like ribosomes are critical intermediates in 40S maturation that link maturation to quality control in tests that mimic ribosome functionality ([33, 34], **Figure 1B**).

To test if the mutation affected the formation of 80S-like ribosomes, we used a previously described assay that takes advantage of the observation that 80S-like ribosomes accumulate to high levels in yeast strains depleted of the essential ATPase Fap7 [20, 21, 33]. Combining Fap7 depletion with mutations that impair the formation of 80S-like ribosomes reduces their accumulation. The formation of 80S-like ribosomes is then measured using sucrose gradient centrifugation, combined with Northern and Western analysis to identify the sedimentation of the 18S rRNA precursor (20S rRNA), and the remaining assembly factors, including Ltv1, respectively. We therefore analyzed lysates from yeast depleted of Fap7 and expressing wt or mutant Ltv1 using sucrose gradients and probed the fractions for 20S rRNA and assembly factors (**Figure 2B&C**). As shown in **Figure 2B**, Ltv1_L216S indeed shifted some of the 20S pre-rRNA from the 80S fraction back to the 40S fraction, demonstrating that the mutations impairs the formation of 80S-like ribosomes.

We and others have previously shown that the formation of 80S-like ribosomes requires the phosphorylation of Ltv1 [20, 29], as well as the ordered release of Ltv1, Enp1 and Rio2 (**Figure 1B**, [21]). Intriguingly, the L216S mutation impairs Ltv1 phosphorylation (**Figure 2C**). Because Ltv1 phosphorylation is required for its release [20, 29], Ltv1_L216S also accumulates in pre-40S ribosomes, with less of it in the free fraction (**Figure 2C&D**). Moreover, as Ltv1 release is required for Enp1 and Rio2 dissociation from pre-40S [21], the Ltv1 mutation also leads to accumulation of Enp1 and Rio2 in pre-40S (**Figure 2C&D**). Thus, together these data indicate that the Ltv1_L216S mutation impairs 40S ribosome maturation, by interfering with the phosphorylation of Ltv1, and its subsequent release, ultimately preventing the formation of 80S-like intermediates.

Phosphomimetic (aspartate) mutations in the Ltv1 phosphorylation site (S336, S339, S342, Ltv1_S/D) can rescue the depletion of the Hrr25 kinase [20], and promote release of Ltv1 in the absence of Hrr25, as well as in mutants that block Ltv1 phosphorylation [21]. We therefore combined the phosphomimetic S/D mutation with L216S to produce Ltv1_L216_S/D and asked if the S/D mutation could rescue Ltv1 release as expected and as seen in all other cases. Surprisingly, Ltv1_L216_S/D remains accumulated in pre-40S ribosomes, and similarly, neither Enp1 nor Rio2 release was rescued (**Figure 2E&F**). Consistently, the S/D mutation exacerbated rather than rescued the growth defect from the Ltv1_L216S mutation (**Figure 2G**).

The observation that Ltv1_L216S cannot be phosphorylated by Hrr25, and that a phosphomimetic mutation cannot rescue the release or growth defects from the L216S mutation, suggest that Ltv1_L216S is positioned differently on pre-40S ribosomes, so that it is no longer accessible to the Hrr25 kinase, and such that S336, S339 and S342 no longer weaken binding when mutated to phosphomimetic aspartate residues. In contrast, the interface with Enp1 appears to remain largely unchanged: an Enp1 mutation, Enp1_WKK (W224V, K228E, K231E), rescues the non-phosphorylatable alanine mutations in Ltv1, Ltv1_S/A (**Figure S2A**), because it leads to weaker Ltv1 binding (**Figure S2B**). Enp1_WKK also rescues Ltv1_L216S (**Figure S2C**), suggesting that the interface between Enp1 and Ltv1 is the same in the Ltv1_S/A and Ltv1_L216S mutants.

Together, the data in this section demonstrate that Ltv1_L216S impairs ribosome assembly, because it cannot be phosphorylated and released, and strongly suggest that these defects arise because Ltv1 is mispositioned on pre-40S ribosomes.

### Ltv1 wraps around the side of the head

To better understand how the Ltv1_L216S mutation alters its function in the context of the genetic and biochemical data that indicate mispositioning of the mutant protein, we wanted to visualize the location of the mutation in the context of the pre-40S structure. Unfortunately, only a small portion of Ltv1 has been visualized in current cryo-EM structures [14, 15]. We therefore obtained a predicted structure for Ltv1 from the alpha-fold website (https://alphafold.ebi.ac.uk/), and then utilized the matchmaker function in Chimera [35] to place this structure on the pre-40S structure using the part of Ltv1 that has been previously visualized as a template.

Overall, this structure for Ltv1 and its placement in pre-40S does not lead to significant steric clashes, but rather is compatible with the pre-40S structure and earlier pre-40S structures (**Figure 3A&Figure S4B**). In the composite structure Ltv1 contacts Rps3, Rps15, and Rps20, which are all validated binding partners for Ltv1 [24, 29–31], thus supporting the placement. Moreover, aspartate 113 and glutamate 115 in Rps20, whose mutation to lysine (Rps20_DE) weakens Ltv1 binding to pre-40S [26], are directly at the Ltv1/Rps20 interface (**Figure 3B)**. Moreover, while Rps20_DE bind directly to K7/K10 in Rps3 (Rps3_KK), their mutation has opposite genetic interactions with mutations in or deletion of Ltv1, suggesting that this interaction was not made when Ltv1 was bound [26]. Indeed, the proposed structure places Ltv1 between those two RPs. Thus, the composite structure is consistent with and explains available data.

**Figure 3:**
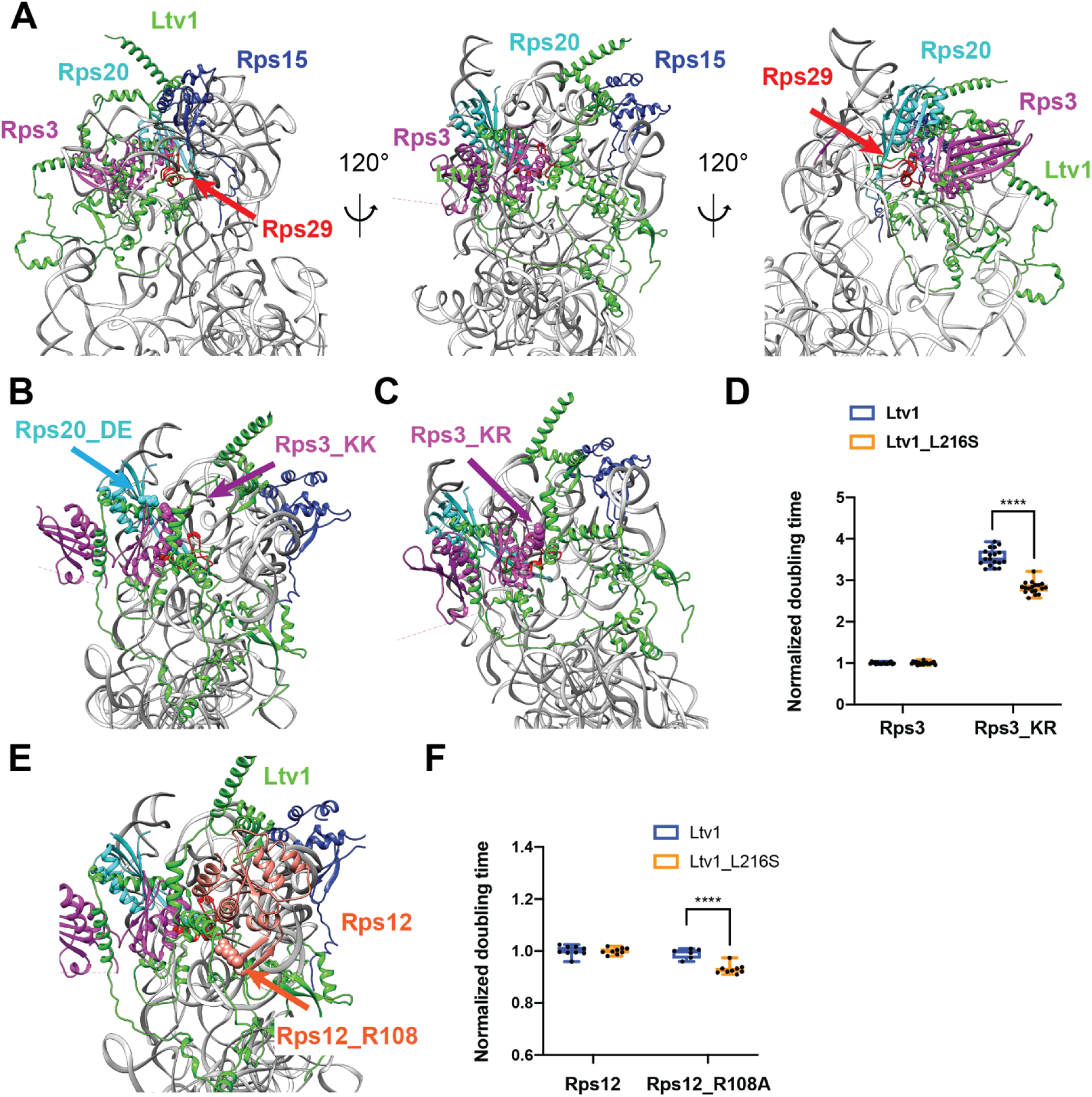
Ltv1 binding to pre-40S ribosomes. (A) A structure for Ltv1 was obtained from the alpha-fold website (https://alphafold.ebi.ac.uk/) and placed on the structure for pre-40S ribosomes (PDB ID 6FAI), by overlay with the Enp1-binding alpha-helix. The composite structure does not produce significant clashes with pre-40S ribosomes and shows interactions between Ltv1 and Rps3, Rps15 and Rps20. All structure displays were generated with Chimera [35]. (B) Detail of the complex demonstrating that Ltv1 separates Rps20_DE from Rps3_KK (highlighted in cyan and purple spheres, respectively), consistent with previous biochemical and genetic data. (C) Detail of the complex demonstrating that Ltv1 binds Rps3K75R76 (Rps3_KR, highlighted in purple spheres). (D) Doubling times (normalized to wt Rps3) for yeast cells expressing either wt Rps3 or Rps3_KR and either wt Ltv1 or Ltv1_L216S. Significance was tested using an unpaired t-test. ****, P<0.0001. (E) Detail of the complex demonstrating that Ltv1 binds Rps12, with a contact from Rps12_R108 (highlighted in red sphere). (F) Doubling times (normalized to wt Rps12) for yeast cells expressing either wt Rps12 or Rps12_R108A and either wt Ltv1 or Ltv1_L216S. Significance was tested using an unpaired t-test. ****, P<0.0001.

To further validate this structure and the placement of Ltv1, we next designed point mutants in ribosomal proteins, which are expected to disrupt Ltv1/RP interactions, and then tested their genetic interaction with Ltv1_L216S.

Inspection of the structure for Ltv1 on the nascent 40S revealed an interaction between Ltv1 and lysine residues K75 and R76 in Rps3 (**Figure 3C**). We therefore tested whether mutation of these residues to glutamates (Rps3_KR) influences cell growth. In fact, Rps3_KR grows over 3-fold more slowly than wt Rps3 (**Figure 3D**), demonstrating the importance of this residue for ribosome assembly and/or function. Next, we combined Rps3_KR with Ltv1_L216S to test for genetic interactions. Notably, Ltv1_L216S and Rps3_KR are epistatic, consistent with the idea that Ltv1 mispositioning in the Ltv1_L216S mutant perturbs the interaction between Ltv1 and Rps3_KR, therefore validating the position of Ltv1 in the composite structure. This observation also suggests that at least part of the growth defect in the Rps3_KR mutant reflects a defect in ribosome assembly.

In addition to interactions with Rps3, Rps15 and Rps20, which have been previously validated [24, 29–31], in the proposed structure Ltv1 also binds Rps12 (**Figure 3E)**. To confirm this interaction and further validate the Ltv1 placement, we tested for genetic interactions between Rps12 and Ltv1. The point of contact between these two proteins includes Rps12_R108. We therefore mutated this residue to alanine (Rps12_R108A). Indeed, when combined with Ltv1_L216S, Rps12_R108A is epistatic (**Figure 3F**), supporting an interaction between the two proteins. Synthetic negative interactions between Rps12 deletion and Ltv1 deletion or Enp1 point mutants have also been previously reported [36] and are consistent with an interaction between these proteins.

Together, these mutational and genetic analyses, combined with the previous biochemical data validate the structure of Ltv1. Additional validation comes from the analysis of rRNA mutations in the next section.

### Ltv1_L216S is located adjacent to helix 34 and perturbs folding of junction j34-35-38

The structure reveals that L216 is part of a short alpha-helical segment adjacent to helix 34 (h34, **Figure 4A**). H34 is part of a 3-helix junction with helices 35 and 38 (j34-35-38). In mature 40S, j34-35-38 forms a tertiary interaction with the helical loop closing helix 31 (l31). We have previously shown that j34-35-38 is misfolded in the absence of Ltv1 [21, 22, 26]. Moreover, our biochemical and previous structural data indicate that j34-35-38 folds once Ltv1 dissociates from pre-40S ribosomes [14, 15, 22]. These observations are consistent with the position of Ltv1 adjacent to h34 revealed here and indicate a direct role for Ltv1 in chaperoning the folding of this rRNA segment.

**Figure 4:**
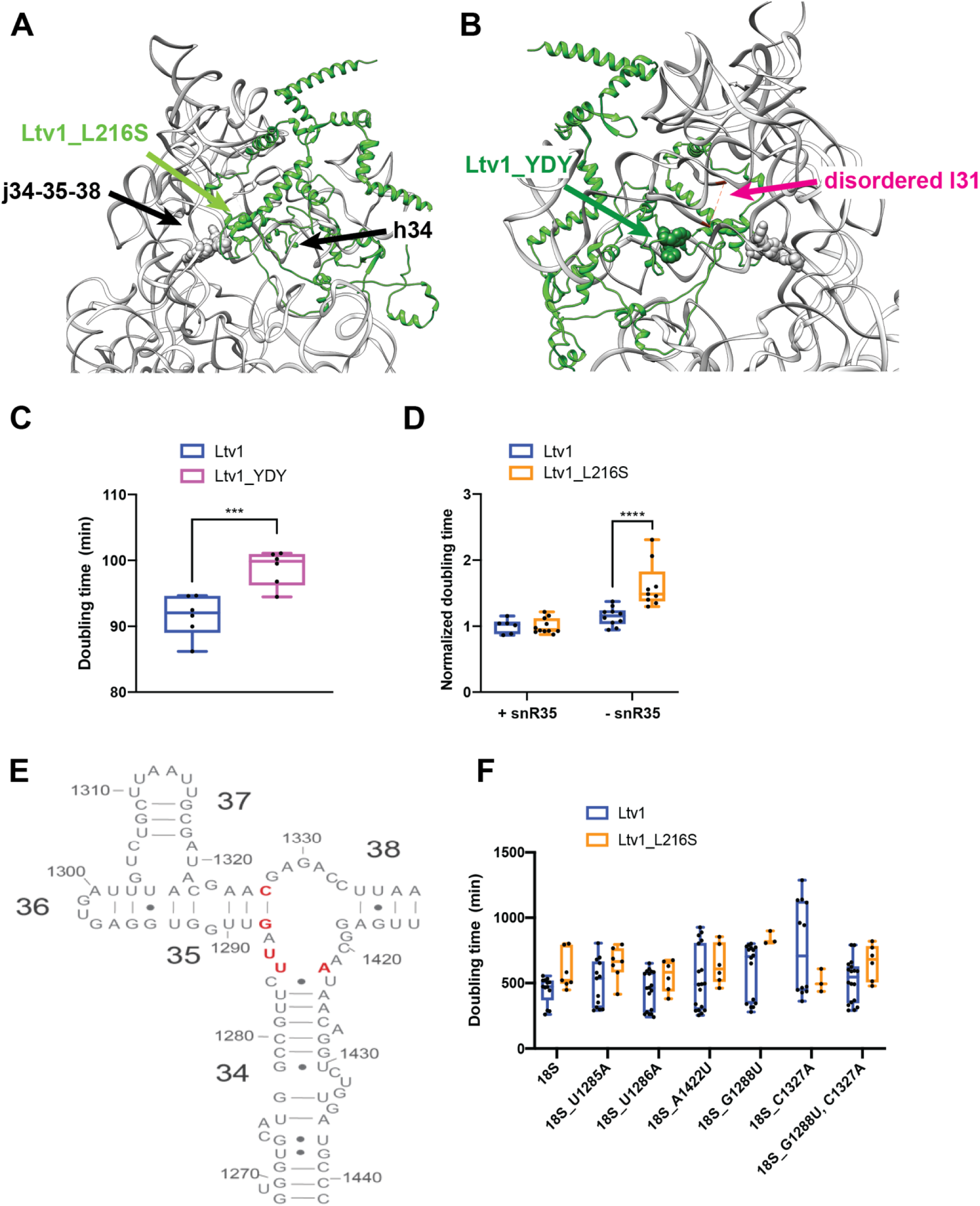
The LIPHAK syndrome associated mutation Ltv1_L216S binds adjacent to j34-35-38 and affects its folding. (A) Detail of the complex demonstrating that Ltv1_L216S (highlighted in green spheres) binds adjacent to j34-35-38 and h34. Residues mutated in panel E are highlighted in gray sphere. (B) Detail of the complex demonstrating that Ltv1_YDY (highlighted in dark green spheres) binds adjacent to l31 (which is disordered in this structure and thus not visible). Its start and end are indicated in magenta and connected by a dashed line. Residues mutated in panel E are highlighted in gray sphere. (C) Doubling for yeast cells expressing either wt Ltv1 or Ltv1_YDY. Significance was tested using an unpaired t-test. ***, P<0.001. (D) Doubling times (normalized to wt snR35) for yeast cells containing or lacking snR35 and expressing either wt Ltv1 or Ltv1_L216S. Significance was tested using an unpaired t-test. ****, P<0.0001. (E) Secondary structure diagram of j34-35-38 illustrating the locations of the mutations in panel F. (F) Doubling times (normalized to wt Ltv1) for yeast cells containing a temperature sensitive RNAPol I mutation and expressing plasmid encoded wt or mutant 18S rRNA and wt Ltv1 or Ltv1_L216S. Significance was tested using an unpaired t-test. ****, P<0.0001.

However, premature folding of l31 leads to misfolding of j34-35-38, as it stabilizes misfolded structures of the junction [22]. The structure shows that Ltv1 also contacts l31 (**Figure 4B**), and mutation of residues Y82, D83, and Y84, which are adjacent to l31 to VAV, to make Ltv1_YDY produces a small growth defect (**Figure 4C**), therefore further validating the structure. However, this interaction between Ltv1 and l31 also opened the possibility that the absence of Ltv1 leads to j34-35-38 misfolding via j34-35-38 directly, or via l31, or both.

To test whether Ltv1 chaperones the folding of j34-35-38 directly, or rather indirectly affects its folding by preventing the premature folding of l31, or both, we carried out a series of genetic experiments. Deletion of snR35, a snoRNA which directs pseudouridylation of U1191 in l31, and prevents the premature folding of l31, results in misfolding of j34-35-38 [22]. This is because the folding of l31 stabilizes misfolded structures in j34-35-38. If a mutation in Ltv1 is epistatic with snR35 deletion, it would suggest that it leads to premature folding of l31 in the same way as snR35 deletion does. In contrast, synergistic effects would indicate that its effects are not via l31. We therefore combined the Ltv1_L216S mutation with the snR35 deletion. The data in **Figure 3D** show that deletion of snR35 is synthetically sick with the Ltv1_L216S mutation, indicating that the effects on j34 misfolding largely arise directly from its location at j34 and not from premature folding of l31.

To further confirm this conclusion, we next tested whether rRNA mutations in the helices that surround the junction and are adjacent to Ltv1 would be sensitive to the Ltv1_L216S mutation. We therefore mutated U1285, U1286 and A1422, which are part of a bulge at the top of helix 34, as well as G1288 and C1327, the base pair which closes h35 (**Figure 4E**), and tested their growth defects alone, and in combination with Ltv1_L216S.

For this purpose, we deleted the Ltv1 gene from a strain where RNA polymerase I has a mutation that renders it temperature sensitive, thereby shutting off all transcription from the endogenous rDNA genes above 32 °C [37]. When supplemented with plasmids encoding either wt rRNA, or a mutation of interest, their impact can then be tested. In addition, we have also supplied either wt Ltv1 or Ltv1_L216S. By themselves the rRNA mutations have small to no effect on growth (data not shown). However, when combined with Ltv1_L216S, they demonstrate strong synthetic sick effects (**Figure 4F**). Together, these data demonstrate a direct role for Ltv1 in enabling the correct folding of j34. Moreover, the observation of j34-35-38 misfolding in the absence of or with Ltv1 mutants further validates the structure, which places Ltv1 directly adjacent to j34-35-38.

### Head protein assembly is globally perturbed in the Ltv1_L216S mutant

Given that the Ltv1_L216S mutant is only partially functional, we wanted to use its defects to gain additional insight into the functions of the wild type protein. Our previous work had shown that Asc1 and Rps10 were substoichiometric in ribosomes lacking Ltv1 [26], suggesting mispositioning of Rps3, their common binding partner. Moreover, genetic experiments also indicated Rps20 was mispositioned or reduced. To investigate RP recruitment to ribosomes globally, we purified 40S ribosomes from yeast expressing either wt Ltv1 or Ltv1_L216S, and then analyzed RP composition via semi-quantitative mass spectrometry. We obtained three independent biological replicates from each strain, normalized the peptide reads for each protein to the reads for the entire sample to normalize for differences in sample abundance, and quantified RP occupancy in the L216S mutant relative to wt Ltv1. These data reveal a striking depletion of most RPs, except Rps3, Asc1, Rps20 and Rps28, from the small ribosomal subunit head in the ribosomes from mutated yeast **(Figure 5A&B**).

**Figure 5:**
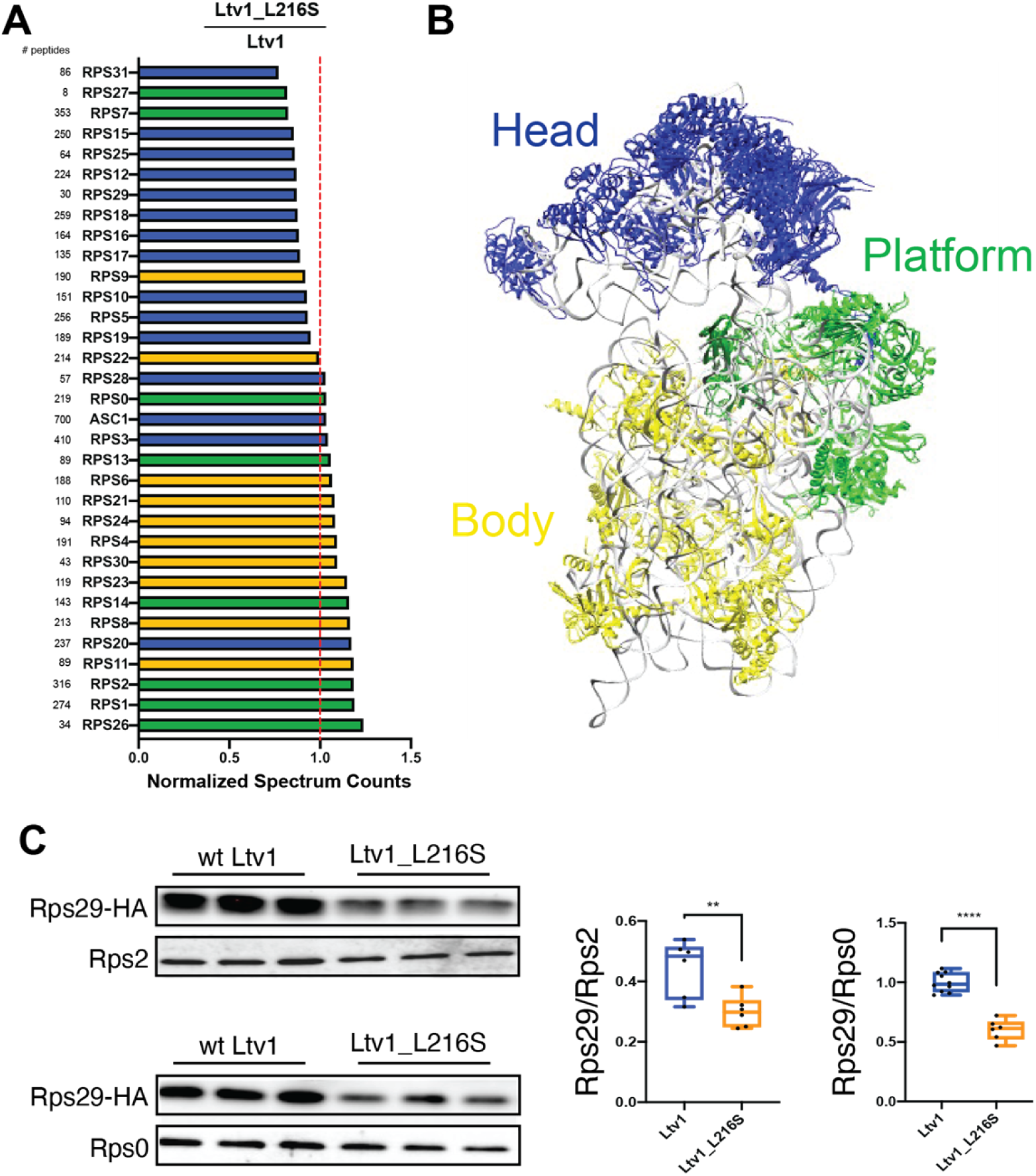
The Ltv1_L216S mutation globally disrupts head assembly. (A) Mature ribosomes from yeast expressing wt Ltv1 or Ltv1_L216S were purified and analyzed by mass spectrometry. Spectral counts for each protein in each sample were normalized by the total number of spectra, and abundance of each protein in Ltv1_L216S ribosomes normalized by the abundance in wt Ltv1 ribosomes. Number of spectra for each protein are indicated on the far left. Ribosomal proteins are color-coded for the substructure to which they bind, with head-binding proteins in blue, platform proteins in green, and proteins from the body in yellow. Averaged data from three biological replicates are shown. (B) Structure of mature 40S ribosomes highlighting the different substructures with the same color-code as in (A). (C) Western analysis of ribosomes purified from cells expressing Rps29-HA and either wt Ltv1 or Ltv1_L216S. Rps29 occupancy was quantified relative to Rps2 and Rps0. Significance was tested using an unpaired t-test. **, P<0.01; ****, P<0.0001.

### DMS-MaPseq reveals global roles for Ltv1 in 40S structure

To better understand the molecular basis for the loss of many head RPs and determine whether additional rRNA regions are misfolded in the ribosomes from the L216S mutant yeast, we utilized DMS-MaPseq to probe the structure of mature 40S subunits from yeast expressing wild type Ltv1 or L216S Ltv1. As previously described [22, 38], ribosomes were exposed to DMS or mock treatment, and after quenching the reaction, 18S rRNA was fragmented, reverse transcribed, and sequencing libraries prepared. After sequence alignment, modified nucleotides are misread in the reverse transcription reaction and appear as a “mutation” relative to the genomic DNA (**Figure S3A**). Thus, the mutational rate at each nucleotide is a measure of its propensity to be modified by DMS. Analysis of the sequencing data indicates high read density over the entire molecule **(Figure S3B**), with some drop off near the 3’-end, due to the methylation of A1781/A1782 (**Figure S3C**). We next compared the mutational rate for each nucleotide in wt and mutant cells, identifying 14 residues, that were more accessible in the mutant and 1 residue that was more protected in the mutant (**Figure 6A-B**, **Table S1)**.

**Figure 6:**
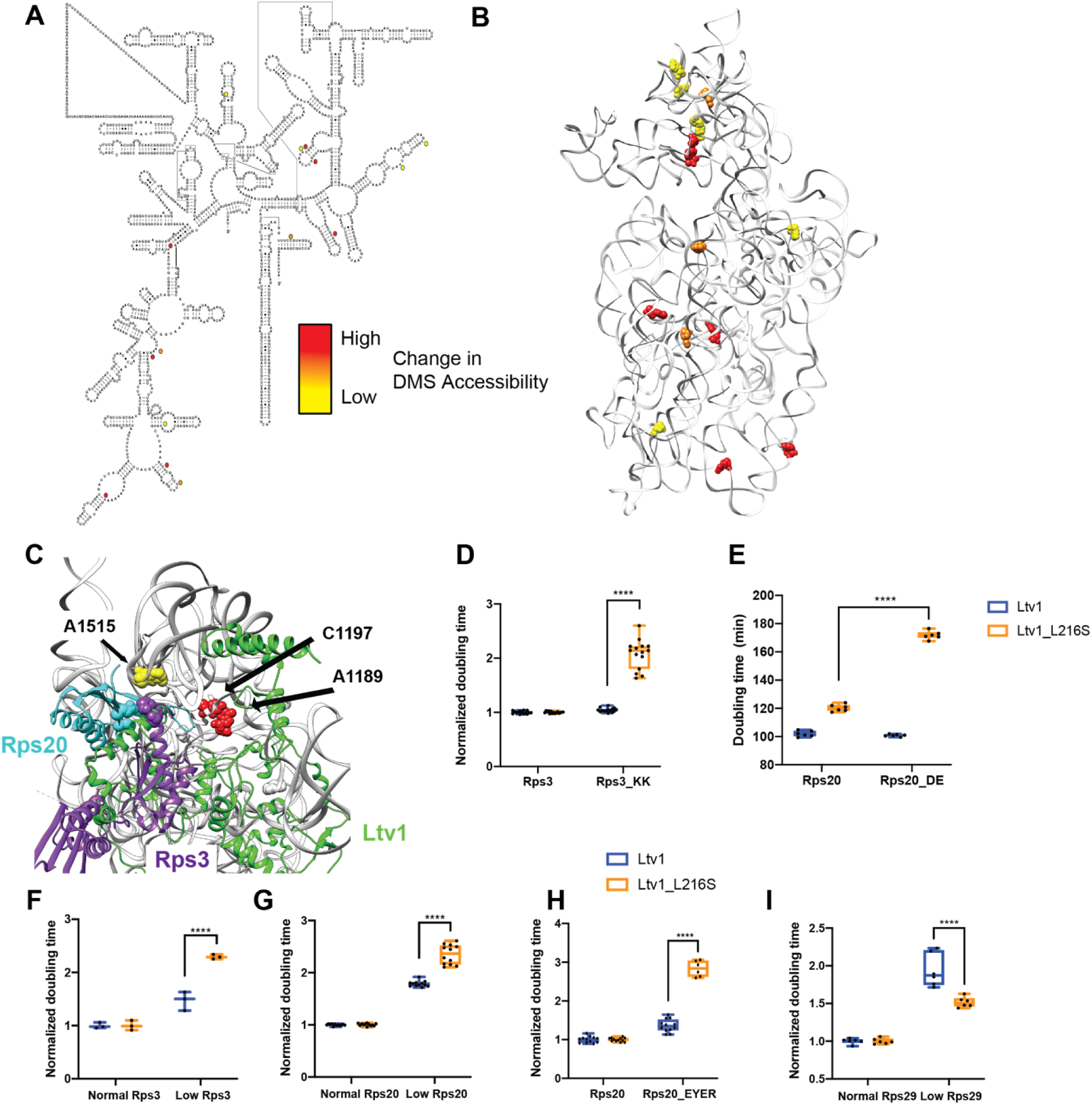
DMS MaPSeq demonstrates perturbations in rRNA folding arising from the Ltv1_L216S mutation. (A) Secondary structure of 18S rRNA demonstrating residues significantly altered in ribosomes from Ltv1_L216S cells. All residues (except C191) are more accessible in ribosomes from Ltv1_L216S. The extent of the alteration is indicated by yellow, orange or red circles. (B) Residues significantly altered in ribosomes from Ltv1_L216S cells mapped onto the 3D structure of 18S rRNA from pre-40S ribosomes (PDB ID 6FAI). Color scheme as in panel A. (C) Highlight of the composite structure from Figure 3, demonstrating that A1515, A1189 and C1197 are adjacent to residues in Ltv1. A1196 is not resolved in the structure and therefore not shown. Rps20_DE and Rps3_KK are shown in cyan and purple spheres, respectively. The Ltv1L216S mutation demonstrates significant synthetic sick or lethal interactions with Rps3_KK (D) or Rps20_DE (E), respectively. (F) Doubling times (normalized to normal Rps3) for yeast cells expressing either normal or low levels of Rps3 and wt Ltv1 or Ltv1_L216S. Normal levels of Rps3 and Rps20 were obtained by using the TEF2 promoter-driven plasmids. Low levels of Rps3 and Rps20 were obtained by using Tet-promoter-driven plasmids and addition of 0 or 20 ng/ml of doxycycline (dox), for Rps3 and Rps20, respectively. Significance was tested using an unpaired t-test. ****, P<0.0001. (G) Doubling times (normalized to normal Rps20) for yeast cells expressing either normal or low levels of Rps20 and wt Ltv1 or Ltv1_L216S. Significance was tested using an unpaired t-test. ****, P<0.0001. (H) Doubling times (normalized to wt Rps20) for yeast cells expressing either wt Rps20 or Rps20_EYER and wt Ltv1 or Ltv1_L216S. Significance was tested using an unpaired t-test. ****, P<0.0001. (I) Doubling times (normalized to normal Rps29) for yeast cells expressing either normal or low levels of Rps29 and wt Ltv1 or Ltv1_L216S. Low levels of Rps29 were obtained by using a Tet-promoter-driven plasmid, which even without dox addition produces reduced levels of Rps29. Normal levels of Rps29 were produced with TEF2 promoter-driven plasmids. Significance was tested using an unpaired t-test. ****, P<0.0001.

Six of the 18S rRNA residues with altered DMS accessibility are located in the 40S head. In addition, there is an additional cluster of 5 residues with altered DMS accessibility on the subunit interface (**Figure 6B**). This cluster will be discussed separately below (*The Ltv1_L216S mutation impairs movement of Tsr1 to the beak*).

### The L216S mutation leads to premature formation of the Rps20-Rps3 contact, which blocks Rps29 binding

Four of the residues that are more DMS accessible in ribosomes from Ltv1_L216S cells, A1189, A1196, C1197 and A1515 are directly adjacent to residues in Ltv1 (**Figure 6C**), further supporting the suggested placement of Ltv1 in pre-40S ribosomes. However, none of them are close to L216, strongly supporting the conclusion above, based on genetic interactions, that Ltv1 is globally mispositioned in the Ltv1_L216S mutant.

Of the residues with altered DMS accessibility, A1515 is directly adjacent to the residues mutated in Rps20_DE and Rps3_KK, which form a contact in mature 40S subunits. Previous data had suggested that Ltv1 blocks the formation of this contact between Rps20 and Rps3 [26], as they have opposite genetic interactions, and because mutation of Rps20 but not Rps3 weakens Ltv1 binding. Consistent with these previous data, in here we show that Ltv1 is placed between these residues, interrupting this contact (**Figure 6C and Figure 3B**). Thus, the increased accessibility of A1515 in the mutant ribosomes indicates that the L216S mutation leads to repositioning of Ltv1 near Rps3 and Rps20. Presumably this would allow for the premature formation of the contact between these two proteins. Consistently, Rps3 and Rps20 are (together with Asc1, which binds directly to Rps3), the only proteins that are *not* depleted from the head of the 40S subunit in ribosomes from Ltv1_L216S cells (**Figure 5A&B**). We therefore hypothesized that the premature formation of the Rps3-Rps20 contact would lead to mispositioning of Rps3/Rps20 and/or impair the recruitment of other RPs.

To test if the premature formation of the Rps3-Rps20 contact would lead to mispositioning of Rps3/Rps20, we combined the Rps20_DE and the Rps3_KK mutations with Ltv1_L216S. Indeed, both Rps3_KK and Rps20_DE demonstrate strong synthetic growth defects with Ltv1_L216S (**Figure 6D&E**). Moreover, reducing the expression of Rps3 or Rps20 using a tet-repressible promoter also is synthetically sick with the Ltv1_L216S mutation (**Figure 6F&G**). Thus, repositioning of Ltv1 due to the L216S mutation to allow for the premature formation of the contact between these two proteins impairs proper positioning of Rps3 and Rps20, rendering them sensitive to depletion and mutation.

To determine whether premature formation of the Rps3/Rps20 contact also impaired the recruitment of other RPs, we considered the structural data illuminating the folding of the head domain rRNA and the recruitment of its RPs [17]. Binding of Rps20 coincides with the appearance of helix 39 (h39) in cryo-EM structures (**Figure S4A**), presumably because Rps20 stabilizes its tertiary contact with h41 in the head. Notably, Rps29 is located not just under Rps20 and Rps3, but its access to its binding site is blocked once h39 docks into place (**Figure S4A**). Thus, Rps29 must bind prior to the docking of h39, and thus prior to Rps20. We therefore considered that the premature formation of the contact between Rps3 and Rps20 would impair the recruitment of Rps29. Indeed, the mass-spectrometry data indicated that Rps29 is one of the most-depleted proteins in the ribosomes from Ltv1_L216S cells (**Figure 5A**). However, Rps29 was detected by relatively few peptides, rendering this measurement unreliable. We therefore used Western analysis with HA-tagged Rps29 to confirm the mass spectrometry data. We constructed a yeast strain where Rps29 is HA-tagged and Ltv1 deleted, supplemented this strain with wt and mutant Ltv1 and purified ribosomes from this strain. Western blotting against the HA-tag on Rps29 demonstrates that indeed Rps29 is reduced by ∼ 50% in ribosomes from Ltv1_L216S yeast (**Figure 5C**).

However, our structure also indicates direct binding interactions between Ltv1 and Rps29 (**Figure 3A**), which likely also contribute to Rps29 recruitment. We therefore carried out additional genetic interaction experiments to obtain data for the ordered binding of Rps29 and Rps20, and the requirement for a concentration-independent step in Rps29 binding. We mutated four residues in Rps20 which form interactions with Rps29: E80, Y82, E83 and R95, thereby producing Rps20_EYER [21]. In addition, we produced a yeast strain, where the Rps29 concentration can be reduced via addition of doxycycline (dox). We combined each alteration with expression of either wt Ltv1 or Ltv1_L216S. Importantly, while Rps20_EYER is synthetically sick with Ltv1_L216S (**Figure 6H**), reducing Rps29 concentration has no effect on Ltv1_L216S yeast, even though it reduces growth in yeast with wt Ltv1, demonstrating strong epistasis (**Figure 6I**). Together, these two observations strongly suggest that the Rps20_EYER mutation rather than affecting Rps29 recruitment, affects Rps20 recruitment, as otherwise the mutation would have the same effect as Rps29 depletion. Consistently, reducing Rps20 with Ltv1_L216S shows the same synthetic sick interaction as Rps20_EYER (**Figure 6G**). Thus, Rps29 must be present before Rps20. Moreover, the epistatic effect from the reduction in Rps29 concentration and Ltv1_L216S is consistent with a concentration-independent step, limiting Rps29 recruitment.

Thus, the genetic interactions, mass spectrometry and DMS-MaPseq data, together with structural analyses indicate that in wt yeast Rps29 binds before Rps20, because Ltv1 blocks the premature formation of the Rps20/Rps3 contact. Mispositioning of Ltv1 via the L216S mutation results in the premature formation of the contact between Rps3 and Rps20. This leads to mispositioning of Rps3 and Rps20, but also blocks Rps29’s access to its binding site, likely due to the premature formation of h39.

### The Ltv1_L216S mutation affects an interaction between the C-terminal tails of Ltv1 and Rps15 with Tsr1

As described above, a second cluster of RNA residues altered in its DMS accessibility in the L216S mutant is on the subunit interface (**Figures 6B&7A&B**). Intriguingly, four of the subunit interface rRNA residues altered in the Ltv1_L216S mutant are located around the binding site for the assembly factor Tsr1 (**Figure 7A**), in particular around an N-terminal helix, which acts as a hinge to allow it to swing towards the beak in 80S-like ribosomes [16], indicating that altered functionality of Tsr1 is responsible for their altered DMS accessibility. On the subunit interface Tsr1 is close to and likely interacts with the C-terminal tails (CTT) of Ltv1 and Rps15 (**Figure S5A**), which could both mediate an effect from the Ltv1_L216S mutation on Tsr1. Because the resolution of the available cryo-EM structures in that area is limited and does not unambiguously identify these three proteins, and because the alpha fold predicted structure shows a different path for yeast Ltv1, we first used genetic interactions to obtain evidence for interactions between Ltv1, Rps15 and Tsr1.

**Figure 7:**
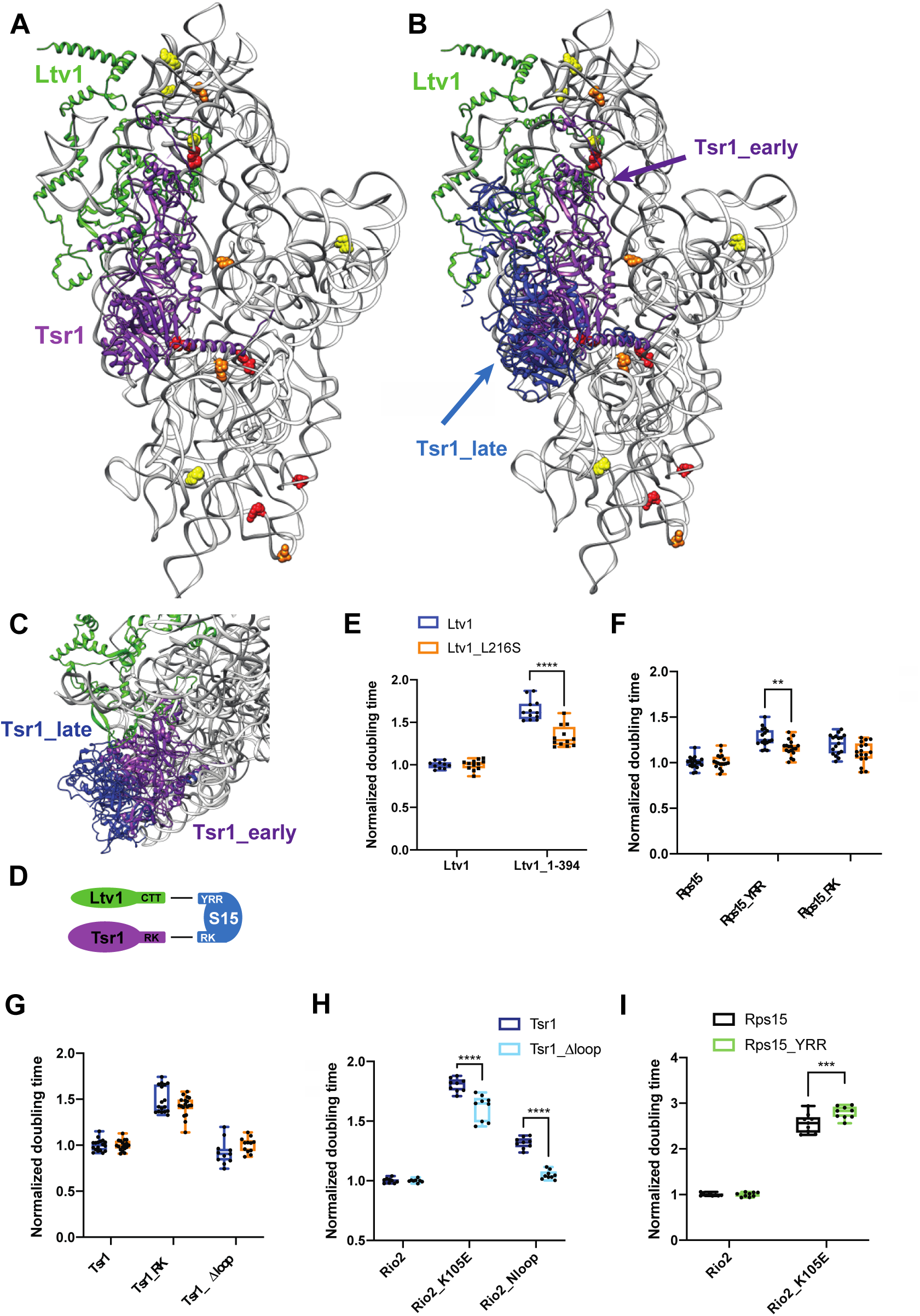
Ltv1 dissociation requires Tsr1 movement to the beak. (A) Structure of the pre-40S ribosome highlighting residues altered in ribosomes from Ltv1_L216S yeast as determined by DMS-MaPseq (Figure 6A&B) showing Tsr1 and Ltv1 (from PDB ID 6FAI). (B) Structure of the pre-40S ribosome with Ltv1 and Tsr1 in the position from earlier pre-40S (6FAI, purple) and from 80S-like ribosomes (in blue), near the beak (6WDR). 6FAI and 6WDR were overlaid by matching the 18S rRNA. (C) Structural detail of the complex in B, viewed from the top of the subunit, demonstrating that Tsr1_early, but not Tsr1_late blocks binding and dissociation of Ltv1. (D) Summary of the genetic interaction network shown by the data in panel E and F and in **Figure S5**. (E) Doubling times (normalized to wt Ltv1) for yeast cells expressing either wt Ltv1 or Ltv1_L216S in full length or truncated (at amino acid 394) form. Significance was tested using an unpaired t-test. ****, P<0.0001. (F) Doubling times (normalized to wt Rps15) for yeast cells expressing either wt Rps15, Rps15_YRR or Rps15_RK and wt Ltv1 or Ltv1_L216S. Significance was tested using an unpaired t-test. **, P<0.01. (G) Doubling times (normalized to wt Tsr1) for yeast cells expressing either wt Tsr1 or Tsr1_RK or Tsr1Δloop and wt Ltv1 or Ltv1_L216S. Significance was tested using an unpaired t-test. (H) Doubling times (normalized to wt Rio2) for yeast cells expressing either wt Rio2, Rio2_K105E or Rio2_loop. Significance was tested using an unpaired t-test. ****, P<0.0001. Data for Tsr1_RK and Tsr1_Δhinge are in Figure S5e of [16] (I) Doubling times (normalized to wt Rio2) for yeast cells expressing either wt Rio2, or Rio2_K105E. Significance was tested using an unpaired t-test. ***, P<0.001. Data for Tsr1_RK and Tsr1_Δhinge are in Figure S5g of [16].

We have previously shown that the growth defect from a mutation in Tsr1, Tsr1_RK (R709E, K712E), which mutates an interaction between Tsr1 and rRNA in the head (**Figure 7B**, see **Figure S5A** for an illustration of the location for mutations in this section), is exacerbated by deletion of the CTT of Ltv1, Ltv1_C394 [16]. Moreover, Tsr1_RK is rescued by a cancer-associated mutation in the CTT of Rps15, Rps15_S136F [21]. These data link Tsr1, Ltv1 and Rps15, and are supported by synthetic negative genetic interactions between Tsr1_RK and Rps15_YRR (Y123I, R127K, R130K [16]). However, because negative genetic interactions are often less reliable and can arise from global perturbations in ribosome assembly, and we had only one rescue, we sought to strengthen the genetic interaction network between Tsr1 and the CTTs of Ltv1 and Rps15.

Notably, we show here that the previously described rescue of Tsr1_RK by an internal deletion in Tsr1, Tsr1_Δloop, requires the CTT of Ltv1 (**Figure S5B**). Moreover, Tsr1_Δloop also rescues the phosphorylation-deficient Ltv1_S/A (**Figure S5C**). Together, these data provide strong evidence for a functional connection between Tsr1 and Ltv1.

Tsr1_RK also demonstrates synthetic genetic interactions with Rps15_YRR, but epistatic interactions with Rps15_RK (**Figure S5D**). These mutations in Rps15 are all in the CTT. In contrast, Ltv1_C394 is neutral to Rps15_YRR, but synthetically sick with Rps15_RK (**Figure S5E**). Together, these data strongly support a functional network between Tsr1 and the CTTs of Ltv1 and Rps15, consistent with the available cryo-EM structures. Moreover, the epistasis data also indicate that the Ltv1_CTT works through the Rps15_YRR residues to affect Rps15, while Tsr1 works through the Rps15_RK residues (**Figure 7D**), the latter being supported by the cryo-EM data (**Figure S5A**).

To decipher if this functional interaction between Ltv1, Rps15 and Tsr1 is disrupted in the Ltv1_L216S mutant, we first combined the Ltv1_L216S mutation with the truncation of its CTT, Ltv1_C394 and measured the growth effects quantitatively. Indeed, these two mutations are epistatic (**Figure 7E**), strongly suggesting that L216S disrupts the interaction between the CTTs of Ltv1 and Rps15. Moreover, the epistatic effect also suggests that the Ltv1 CTT is mispositioned in the L216S mutant. We also tested for genetic interactions between Ltv1_L216S and Rps15_RK and Rps15_YRR and observed epistasis with Rps15_YRR (**Figure 7F**), similar to the epistasis observed with the CTT-deletion of Ltv1 (**Figure 7E**). This finding further supports the conclusions above that (i) the Ltv1 CTT interacts with Rps15 via the YRR residues and (ii) that the CTT of Ltv1 is mispositioned in the L216S mutant. In contrast, both Tsr1_RK and Tsr1_Δloop as well as Rps15_RK are neutral to Ltv1_L216S (**Figure 7G**), indicating that the Ltv1_L216S mutation does not affect the interaction between Tsr1 and Rps15.

Thus, these genetic data demonstrate functional interactions between the CTTs of Ltv1, Rps15 and Tsr1, consistent with the cryo-EM structures. The DMS-MaPSeq data show that residues around the Tsr1 binding site are altered in ribosomes from cells with the LIPHAK mutation, indicating an alteration in this network during assembly. Intriguingly, many of the RNA residues with altered DMS accessibility in the L216S mutant are located at or near the hinge in Tsr1 that enables its movement from the decoding site helix to the beak as 80S-like ribosomes are formed [16]. Thus, the LIPHAK mutation seems to affect the movement of Tsr1 from the decoding site to the beak.

### Movement of Tsr1 to the beak is required for Ltv1 release

We have previously shown that Ltv1 phosphorylation is required for its dissociation [20, 21], but temporally separated, because it also required the phosphorylation of Rio2, as well as another, undefined step (**Figure 1D**, [21]). Importantly, our previous data showed that this step was blocked by mutations in Tsr1 (Tsr1_RK) and Rio2 (Rio2_loop, Rio2_K105E). Moreover, deletion of a long internal loop in Tsr1 (Tsr1_Δloop) rescues the Tsr1_RK mutation [21].

Many of the RNA residues of altered accessibility in the L216S mutant are located at or near a hinge in Tsr1, which allows it to swing away from the decoding site helix towards the beak, when pre-40S are joined by 60S subunits to form 80S-like ribosomes (**Figure 7A-C**, [16]). Importantly, given the location of the N-terminus of Ltv1 *under* Tsr1, this movement of Tsr1 is likely required not just for the formation of 80S-like ribosomes, but also the dissociation of Ltv1.

We therefore hypothesized that the unidentified step in the dissociation of Ltv1 was the hinge-driven movement of Tsr1 to the beak, and that this step was impaired in the Tsr1_RK, the Rio2_loop and Rio2_K105E mutants. To test this model, we wanted to determine if deleting the hinge in Tsr1 also blocked Ltv1 dissociation. Normally, this experiment would involve assessing the binding of Ltv1, Enp1 and Rio2 to pre-40S, and investigating Ltv1’s phosphorylation status [21]. However, because Tsr1_Δhinge binds ribosomes weakly ([16] & data not shown), all other AFs are also partially dissociated, as previously seen [16], and we were unable to directly assess Ltv1 dissociation in sucrose gradients. Instead, we carried out an extensive genetic analysis, to demonstrate that deleting the hinge, which blocks Tsr1’s movement [16], affects the same step affected by Tsr1_RK, Rio2_loop and Rio2_K105E, which is the dissociation of Ltv1 [21].

As described above, Tsr1_Δloop rescues Tsr1_RK [21], as well as Tsr1_Δhinge [16]. Thus, we tested if Tsr1_Δloop also rescued Rio2_loop and Rio2_K105E. Indeed, both mutations are rescued by this deletion in the Tsr1 internal loop, Tsr1_Δloop (**Figure 7H**). Importantly, Tsr1_Δloop also rescues mutation of the Ltv1 phosphorylation site, Ltv1_S/A (**Figure S5C**), demonstrating that it is linked to release of Ltv1. Moreover, Tsr1_RK, Tsr1_Δhinge and Rio2_K105E are all sick with Rps15_YRR (**Figure 7I** and [16], Rio2_loop was not tested). Thus, Tsr1_RK, Tsr1_Δhinge, Rio2_K105E and Rio2_loop have the same genetic interactions. We have previously shown that Tsr1_Δhinge blocks movement of Tsr1 to the beak [16]. Additionally, we have shown that Tsr1_RK, Rio2_loop and Rio2_K105E affect a yet unidentified step to activate the release of phosphorylated Ltv1 ([21], **Figure 1B**). Thus, taken together, these data suggest that the unidentified step that activates the release of Ltv1 is the movement of Tsr1 to the beak, as predicted by the structural data that indicate that Ltv1 is located under Tsr1.

### Blocking movement of Tsr1 to the beak impairs Rps31 recruitment

As described above, in ribosomes from cells expressing the Ltv1_L216S mutant most RPs are depleted from the 40S subunit head (**Figure 5**). Notably the protein that is most significantly depleted is Rps31. Rps31 is encoded as an N-terminal ubiquitin fusion protein, which is posttranslationally cleaved into ubiquitin and Rps31. Previous data strongly suggest that it is incorporated into ribosomes as the fusion protein, as the unprocessed protein is found in polysomes when cleavage is delayed by a point mutation at the processing site [39]. Rps31 is a late-binding RP, which is not yet incorporated in the stable cytoplasmic 40S intermediates that have been visualized by cryo-EM [14, 15], although cryo-EM structures of later 80S-like ribosomes show it present ([16], **Figure 1A**). This is consistent with data demonstrating that depletion of Rps31 leads to accumulation of the cytoplasmic 20S rRNA [39]. Similarly, impairing the removal of the ubiquitin also delays Nob1-dependent cleavage in 80S-like ribosomes [39], as does deletion of the N-terminal extension of the protein [40]. Thus, the available structural and biochemical data suggest strongly that Rps31 is incorporated around the transition between 40S and 80S-like ribosomes.^1^

Intriguingly, if Tsr1 were still located in its stable position near the decoding site, then the N-terminal extension of Rps31 would have to slide under Tsr1 **(Figure 8A&B**). Moreover, in that location, the head group of Tsr1 would clash with the N-terminal ubiquitin fusion (**Figure 8A&B**). Thus, the structures of pre-40S ribosomes suggest that Tsr1 blocks the recruitment of Rps31 to its binding site at the subunit interface, explaining why Rps31 recruitment is delayed until the formation of 80S-like ribosomes, which is accompanied by the hinge-motion-driven movement of Tsr1 ([16], **Figure 8A&B**). Together with the DMS-MaPseq data which demonstrate alterations around the Tsr1 hinge, these data thus suggest that the movement of Tsr1 to the beak is required for Rps31 recruitment.

**Figure 8:**
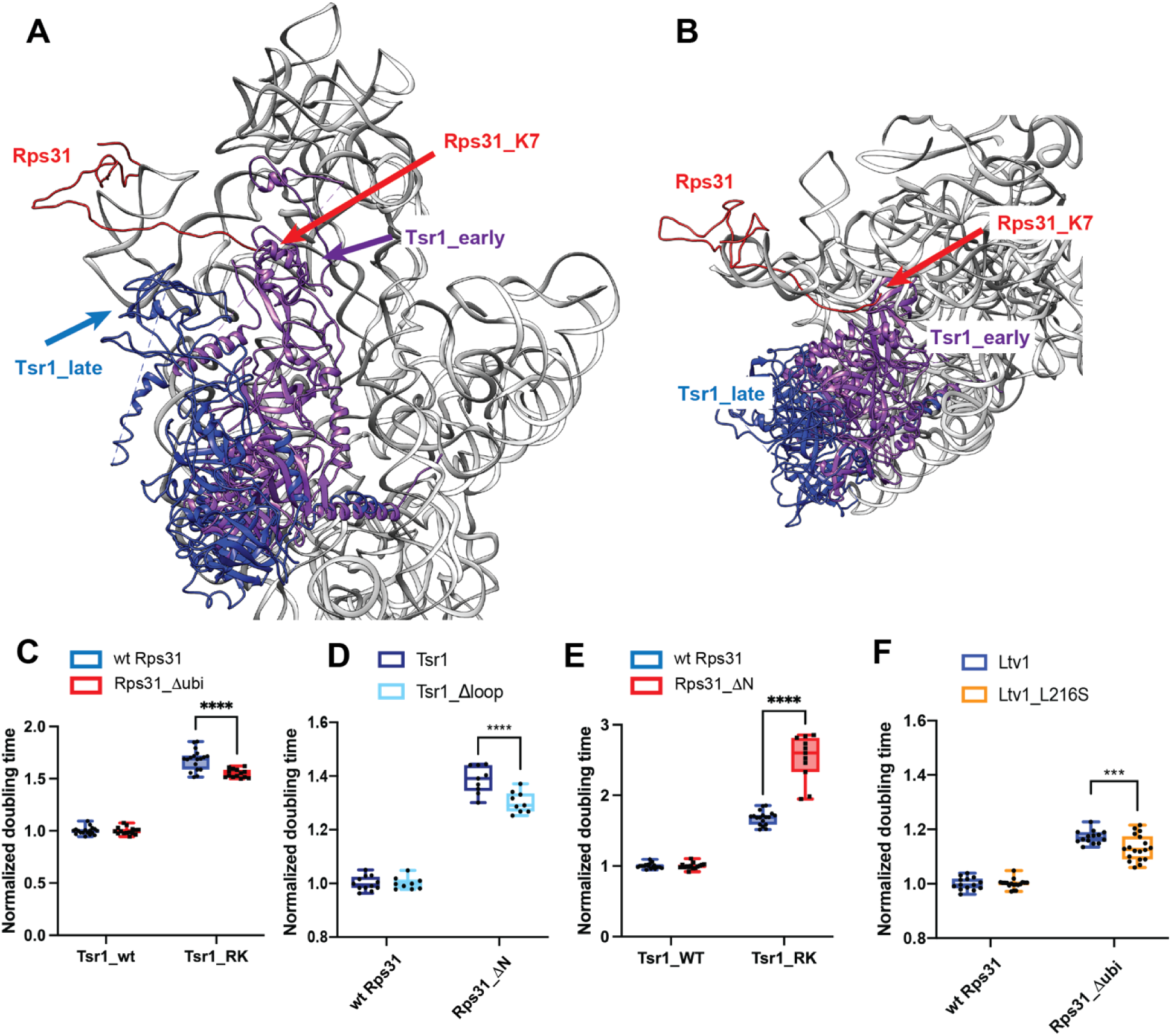
Rps31 binding requires Tsr1 movement to the beak. (A) Composite structure of pre-40S ribosomes (6FAI) with Tsr1 in the early (6FAI) and late position (6WDR) and showing Rps31 from mature ribosomes (3JAM). 6FAI and 3JAM ribosomes were overlaid by matching Rps18, and 6WDR was overlaid onto these by matching 18S rRNA. (B) Structural detail of a top view of the structure in panel A highlighting the N-terminus of Rps31 near Tsr1 in the early position. Note that residues 1-6 are not resolved, the first residue visible is K7. (C) Doubling times (normalized to wt Tsr1) for yeast cells expressing either wt Rps31 or Rps31 lacking the N-terminal ubiquitin fusion (Rps31Δubi), and either wt Tsr1, or Tsr1_RK. (D) Doubling times (normalized to wt Rps31) for yeast cells expressing either wt Rps31 or Rps31 lacking the N-terminal extension (Rps31ΔN), and either wt Tsr1, or Tsr1_Δloop. (E) Doubling times (normalized to wt Tsr1) for yeast cells expressing either wt Rps31 or Rps31 lacking the N-terminal extension (Rps31ΔN) and either wt Tsr1, or Tsr1_RK. (F) Doubling times (normalized to wt Rps31) for yeast cells expressing either wt Rps31 or Rps31Δubi, and either wt Ltv1, or Ltv1_L216S. Significance was tested using an unpaired t-test. ****, P<0.0001.

To test this model, we investigated genetic interactions between Rps31 and Tsr1. If Tsr1_RK and Tsr1_Δhinge impair the movement of Tsr1 to the beak, and this movement is important for the incorporation of Rps31, then we expect that Tsr1_RK is rescued by Rps31_Δubi. Indeed, Tsr1_RK shows small epistatic interactions with Rps31_Δubi (**Figure 8C**).

In addition, previous work has indicated that removing an N-terminal tail in Rps31, which is eukaryote-specific and not conserved in archaea, which also do not conserve Tsr1, is required for assembly of Rps31 into ribosomes [40]. This tail stretches across the front of the head. Given that our data indicate a steric block for Rps31, we combined an Rps31 deletion mutant, lacking this stretch (but containing the ubiquitin), called Rps31_ΔN, with mutants in Tsr1. As seen before [40], Rps31_ΔN produces a strong growth defect. Whether this effect arises because now the ubiquitin is hindered even more by Tsr1, or because this stretch is required for an initial encounter complex, or both, remains unknown. Importantly, the growth defect from Rps31_ΔN is epistatic to Tsr1_Δloop (**Figure 8D)**, with a small but not statistically significant growth rescue. In contrast, Rps31_ΔN is synthetically sick with Tsr1_RK (**Figure 8E)**.

To further support for the model that Ltv1 affects Rps31 incorporation via Tsr1, we combined the Ltv1_L216S mutation with deletion of the ubiquitin in Rps31, Rps31_Δubi. If Ltv1_L216S blocks assembly upstream of the movement of Tsr1 to the beak, then removing the ubiquitin-portion of the gene, which cannot be accommodated without Tsr1 movement, should partially rescue the defects from the L216S mutation. Indeed, while the ubiquitin deletion has no effect on growth in the background of wt Ltv1, it slightly but reproducibly rescues the growth defect of the Ltv1_L216S mutation (**Figure 8F**).

Together, these data provide strong evidence that indeed movement of Tsr1 towards the beak is required for the incorporation of Rps31, and that this movement is impaired by the Ltv1_L216S mutation, thereby explaining the reduced incorporation of Rps31.

## Discussion

### Ltv1 Orchestrates the Hierarchy of Head RP Binding

Taking advantage of a novel mutation in Ltv1, Ltv1_L216S, which is associated with a human dermatological condition known as LIPHAK disease [32], we have uncovered here sweeping roles for Ltv1 in orchestrating the assembly of the small ribosomal subunit head, using a number of different strategies as discussed below.

Previous work has shown that Ltv1 directly binds Rps3, Rps15 and Rps20, and is required for the efficient recruitment of Asc1 and Rps10 [24, 26, 29–31]. Moreover, previous data suggested that Rps3 positioning is affected directly by Ltv1, thereby mediating positioning of Rps17 and Asc1 [26]. Finally, roles in quality control of head assembly have also been described, in particular proofreading of the positioning for Rps3, Rps15, and Rps20 [21]. Here, we expand these roles to demonstrate that Ltv1 also directly binds Rps12 and Rps29 and orchestrates the recruitment of Rps12, Rps29 and Rps31. For Rps12 and Rps29 this is consistent with previous genetic data [26, 36]. Moreover, our data also indicate that correct positioning of Rps15 depends on Ltv1. Thus, Ltv1 directly binds 5 of the 15 head binding RPs (**Figure 9A**) and is involved in the assembly of 9 of them (**Figure 9B)**, thereby explaining the global defects in head assembly that we observe here in ribosomes from cells containing the LIPHAK mutation.

**Figure 9:**
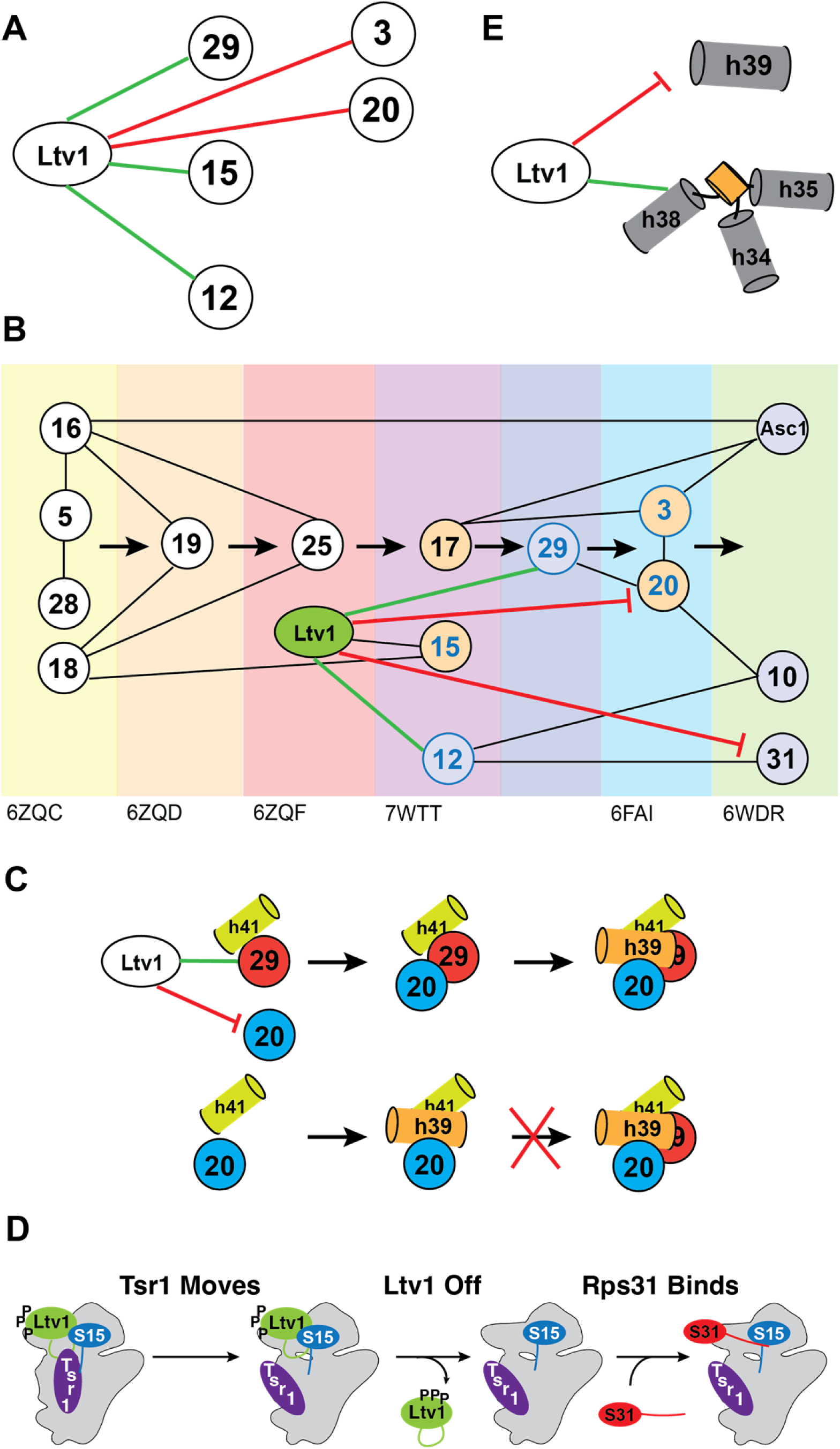
Role of Ltv1 in head assembly. (A) Direct binding interactions for Ltv1. (B) Ltv1 orchestrates the hierarchy of head assembly. Ltv1 is recruited around the same time as Rps25 binds, as determined by the appearance of the C-terminal extension in human ribosomes. RPs that bind directly to Ltv1 are shown in blue circles. RPs Ltv1 helps recruit are connected by a green line and proteins delayed by Ltv1 are connected by a red line. Proteins that are reduced or mispositioned when Ltv1 is absent or mutated are shown in blue or orange spacefill. (C) Ltv1 orchestrates the order of binding of Rps20 and Rps29. By recruiting Rps29 and blocking. The binding of Rps20, Ltv1 enables Rps29 to bind before Rps20, whose binding stabilizes helix 39 (h39) in its tertiary contact with helix 41 (h41). Without Ltv1 (or with the Ltv1_L216S mutation) Rps20 binds first, thus stabilizing h39 and blocking access for Rps29. (D) Tsr1 movement to the beak is required for Ltv1 dissociation and Rps31 binding. Parts of Ltv1 (illustrated by the green loop) bind under Tsr1, requiring its movement for dissociation. This also allows for recruitment of Rps31. Tsr1 movement is regulated via Rps15 and Ltv1. (E) Ltv1 delays formation of h39 and facilitates the formation of j34-35-38.

More importantly, the data herein demonstrate for the first time how Ltv1 carries out its roles. By directly binding Rps12 and Rps29, Ltv1 helps in their recruitment. Moreover, the direct interaction with Rps3, Rps15 and Rps20 enables proper positioning of these RPs, as shown previously for Rps3 and here for Rps15 and Rps20. More surprisingly, the data also demonstrate that by binding to Rps3 and Rps20 directly, Ltv1 prevents the premature formation of their contact, thereby explaining previous genetic and biochemical data [26]. The data in here show that the LIPHAK mutation mispositions Ltv1, thereby dislodging the protein from the Rps20 contact and allowing for the premature formation of the Rps3/Rps20 contact. Importantly, this appears to also misposition Rps3 and Rps20, as the LIPHAK cells are hyper-sensitive to mutations in the Rps3/Rps20 contact. Moreover, we have previously shown that in cells lacking Ltv1 entirely, their interaction partners Rps10 and Asc1 are substoichiometric [26]. Thus, Ltv1 uses its direct binding interactions with 5 head RPs to order their assembly, both by prioritizing some and delaying others.

In addition to using direct binding interactions to orchestrate the RPs assembly hierarchy of the head, Ltv1 also affects recruitment of two RPs indirectly via conformational transitions in the nascent 40S subunit. Our analysis of previous structures of assembling ribosomes indicates that Rps20 binding to nascent ribosomes also coincides with the appearance of h39 in cryo-EM maps, likely because Rps20 binding stabilizes its docking into a tertiary contact with h41 (**Figure S4A)**. Importantly, docking of h39 blocks access of Rps29 into its binding site. Thus, Rps29 must bind before Rps20 enables the docking of h39 (**Figure 9C**). The premature formation of the Rps3/Rps20 contact will therefore prematurely enable the binding of Rps20 and the stabilization of h39 in its tertiary contact and is therefore expected to impair the recruitment of Rps29, which we confirm using ribosome purification and genetic analyses. Thus, Ltv1 enables the recruitment of Rps29 by delaying the binding of Rps20 and the formation of rRNA tertiary structure, thereby keeping open the entry for Rps31.

In addition, our data also demonstrate how Ltv1 orchestrates the recruitment of Rps31. Our genetic data strongly suggest that Rps31 binding is blocked by Tsr1, which has to rotate towards the beak to enable Rps31 binding and Ltv1 dissociation (**Figure 9D)**. This finding is consistent with existing cryo-EM structures that show Rps31 binding occurs with the Tsr1 movement to the beak. Our data show that Ltv1 enables this rotation via contacts between its C-terminal tail, the C-terminal tail of Rps15 and Tsr1 itself. Moreover, our data also strongly suggest that this movement of Tsr1 towards the beak is required for Ltv1 release, recognizing it as a previously identified activation step for Ltv1 release [21].

Ltv1 dissociation is required for Enp1 binding [21], which is explained by the observation that Enp1 is buried under Ltv1 [14, 15]. Moreover, cryo-EM structures demonstrate that Enp1 blocks the binding of Rps10 at its mature site [25]. Thus, Rps10 binding requires Enp1 dissociation, which itself demands Ltv1 release. Thus, Ltv1 also indirectly regulates the timing of Rps10 incorporation.

These observations are summarized in **Figure 9B**, which illustrates the central role of Ltv1 in head assembly and explains much of the previously unexplained parts of the assembly hierarchy. Close inspection of available cryo-EM structures indicates that Ltv1 is recruited to rRNA posttranscriptionally around the time of Rps25 recruitment and after binding of Rps19, as its C-terminal helix at the subunit interface becomes visible then [41]. Ltv1’s binding interactions with Rps15 and Rps12 explain how these are then recruited in the next assembly step, even though their RP interaction partners have been unchanged. The next available structure shows recruitment of Rps3, Rps20 and Rps29, but as described above, the data in here show that Rps29 is recruited first, both via directly binding to Ltv1, and because Ltv1 delays Rps20 recruitment, consistent with structures that show Rps29 being buried once Rps20 is bound and its interacting helix, h39, is docked. Future experiments will be required to dissect how Ltv1 and/or Rps20 are moving to enable the formation of the contact between Rps20 and Rps3, which is required for Ltv1 phosphorylation and release [21]. Finally, the data also explain why Rps31 binding is delayed, even though its only interaction partner, Rps12, is bound earlier.

### Ltv1 Facilitates rRNA Folding

In addition, our data also indicate that Ltv1 plays direct roles in chaperoning the folding of j34-35-38 (**Figure 9E**). We have previously shown that this 3-helix junction folds upon Ltv1 dissociation, indicating a role for Ltv1 in delaying its folding [14, 15, 21, 22]. The data in here strongly suggest that Ltv1 binds this junction directly and demonstrate that this interaction is required for proper folding of j34-35-38. While future experiments will be required to better understand how this (and other) 3-helix junctions fold, and how Ltv1 blocks the formation of its structure, the observation that disrupting h34 and h35, which lead into the junction, exacerbates effects from the LIPHAK mutation indicates that Ltv1 does not disrupt these helices. Instead Ltv1 may bind the connecting elements. Indeed, overlaying an earlier 40S structure indicates that h34 is formed prior to h35 and h38, but not yet positioned correctly (**Figure S4B**). Interestingly, one of the strands of h35 binds directly to Ltv1, suggesting that Ltv1 might function in folding of h34-35-38 either by positioning h35 or delaying the formation of this short helix that emanates from the junction. Future experiments will address how folding of 3 helix junctions can be facilitated.

We have previously shown that Rio2 plays similar roles in delaying the formation of a tertiary contact between the loop of h31 and j34-35-38 [22]. Together, these findings indicate the importance of a carefully orchestrated rRNA folding order to enable efficient ribosome assembly. While Rio2 appears to directly block the formation of a tertiary contact, Ltv1 appears to do this indirectly, by blocking the assembly of an RP, which stabilizes this tertiary contact.

In addition to chaperoning the folding of j34-35-38, our data also strongly suggest roles for Ltv1 in folding of h39, and its tertiary contact with h41 (**Figure 9E**). The structure in here, together with previous biochemical data indicate that Ltv1 blocks the formation of the Rps3-Rps20 contact, thereby delaying the recruitment of Rps20, which binds and stabilizes the docking of h39 into its tertiary interaction. Our previous observation that residues at the tip of h39 are more protected in ribosomes from cells lacking Ltv1 [26], indicating their misfolding, suggest that by delaying this contact between h39 and h41, Ltv1 also chaperones the folding of this helix.

### The Cancer-Associated Rps15-S136F Mutation Enables Unregulated Tsr1 Rotation

We have previously shown that the Tsr1_RK mutation can be bypassed by a mutation in Rps15 that is associated with chronic lymphocytic leukemia (CLL), Rps15_S136F, which leads to defects in the resulting ribosome population [21]. However, it was unclear what step exactly was blocked by the Tsr1_RK mutation. The data in here strongly suggest that the step blocked by the Tsr1_RK mutation is the rotation of Tsr1 towards the beak. Interestingly, Rps15_S136F does not rescue Tsr1_Δhinge (data not shown), indicating that the Rps15 mutation does not affect the movement *per se* but its regulation via the Ltv1/Rps15/Tsr1 network. Given that Tsr1_Δloop rescues all the mutants that block the hinge movement, we speculate that the loop that is removed by the Tsr1_Δloop mutation holds Tsr1 near the beak, via interactions with Rps15, as shown in **Figure S5A**. Thus, the cancer-associated Rps15_S136F mutation, which leads to increased misinitiation at non-AUG codons, may perturb Rps15 directly, or affect how Ltv1 (or other, yet undescribed factors) regulates the interaction between Rps15 and the loop in Tsr1.

## Materials and Methods

### Yeast Strains and Plasmids

*Saccharomyces cerevisiae* strains (**Table S2**) were either obtained from the Horizon Dharmacon Yeast Knockout Collection, or created by homologous recombination [42] and confirmed by serial dilution, PCR, and Western blotting in cases where antibodies were available. Plasmids were generated by standard cloning techniques and confirmed via Sanger sequencing, and are listed in **Table S3**.

### In Vivo Subunit Joining Assay

Gal:Fap7 cells (with the appropriate additional genomic alterations) were grown at 30°C for >16h in YPD to deplete Fap7, and harvested in the presence of cycloheximide as described [21, 33, 43]. Cells were lysed under liquid N_2_, cell debris spun out and the supernatant loaded onto a sucrose gradient as described below.

### Sucrose-Gradient Fractionation

5,000-7,500 OD of clarified lysate (∼ 200 µl) were loaded onto a 10-50% sucrose-gradient and centrifuged at 40,000 rpm for 2 hours using a SW41Ti rotor before fractionation into 700 µl fractions. Northern blot samples were prepared by phenol chloroform extraction on 200 µl of each fraction. For Western blotting, fractions were mixed with SDS loading buffer, and ran on an SDS-PAGE gel, transferred and probed with the appropriate antibodies, as explained in [21, 33, 43].

### Northern Blot

Northern blots were prepared from ΔLtv1 strain supplemented with either Ltv1, or Ltv1_L216S. The cells were grown overnight in minimal media, diluted into fresh YPD and harvested. Total RNA was isolated through phenol/chloroform RNA extraction, denatured in formaldehyde, and separated on an agarose gel. The following probes were used: 20S: GCTCTCATGCTCTTGCC; 18S: CATGGCTTAATCTTTGAGAC; 25S: GCCCGTTCCCTTGGCTGTG; and U2: CAGATACTACACTTG.

### Growth Curve Measurements

Cells were grown in glucose minimal media or YPD overnight, diluted into fresh minimal media or YPD for 3-6 hours, before diluting them into a 96-well plate at a starting OD of 0.04-0.1 and placed in Synergy.2 plate reader (BioTek) to record OD_600_ for 48 hours while the plate was shaking at 30°C.

rRNA mutants were grown in minimal media or YPD at 30°C overnight and diluted into fresh YPD to incubate for 3h at 30°C before changing the temperature to 37°C for 3h. Cells were then inoculated into YPD in a 96-well plate, where the growth was recorded for 48 hours at 37°C.

### Ribosome Purification

Cells were resuspended in ribosome lysis buffer (20 mM Hepes/KOH at pH 7.4, 100 mM KOAc, and 2.5 mM Mg(OAc)_2_) supplemented with 1 mg/ml heparin, 1 mM benzamidine, 2 mM DTT, and protease inhibitor cocktail (Sigma-Aldrich) and lysed under liquid N_2_. The cell lysates were thawed and clarified before layering 200 µl onto 500 µl sucrose cushion and centrifuging at 70k rpm for 65 min in a Ti110 rotor. The supernatant was discarded, and the pellet containing the ribosomes was resuspended in high salt buffer (ribosome buffer, 500 mM KCl, 1 mg/ml heparin, and 2 mM DTT) and layered onto another 500 µl sucrose cushion. The tubes were centrifuged at 100k rpm for 75 min, and the pelleted ribosomes were resuspended in subunit separation buffer (50 mM Hepes/KOH at pH 7.4, 500 mM KCl, 2 mM MgCl_2_, 2 mM DTT and 1 mM puromycin (Sigma-Aldrich), incubated for 1h at 37 °C, and loaded onto a 5-20% sucrose gradient (50 mM Hepes/KOH, pH 7.4, 500 mM KCl, 5 mM MgCl_2_, 2 mM DTT and 0.1 mM EDTA) and centrifuged for 8 h at 30k rpm. To isolate the separated subunits the gradients were fractionated and 40S subunits collected, buffer exchanged into ribosome storage buffer (ribosome buffer with 250 mM sucrose and 2 mM DTT) and concentrated to be flash frozen and stored at - 80°C for subsequent mass spectrometry or Western analysis.

### Mass Spectrometry

Purified ribosomes were prepared as stated above and precipitated in acetone. Three biological replicates of WT Ltv1 and Ltv1_L216S were analyzed.

### DMS MaPseq Sample Preparations

Prior to flash freezing the purified ribosomes were treated with or without 1% DMS (Sigma-Aldrich) in the presence of 80 mM Hepes pH 7.4, 50 mM NaCl, 5 mM Mg(OAc)_2_, and 0.2 µM RNaseP RNA at 30 °C for 5 min. DMS reactions were stopped by the addition of 0.4 volumes stop buffer (1M β-ME, 1.5M NaOAc, pH 5.2), and purified using phenol chloroform precipitation.

### DMS MaPseq RNA Library Preparation

RNA library preparation was adapted from [44] and performed as described [22, 38]. Briefly, RNAs were fragmented with the addition of 20 mM MgCl_2_ at 94 °C for 10 min. Fragments were separated on a denaturing 15% gel, and fragments between 50-80 nt were cut out and RNA was eluted overnight in RNA gel extraction buffer (0.3 M NaOAc, pH = 5.5, 1 mM EDTA (ethylenediaminetetraacetic acid), 0.25% vol/vol SDS (sodium dodecyl sulfate). Eluted RNA fragments were dephosphorylated using T4 PNK (NEB), ligated to an adaptor using T4 RNA ligase II truncated KQ(NEB), and then gel purified. TGIRT III (InGex) was used to reverse transcribe RNAs in 50 mM Tris HCl, pH = 8.3, 75 mM KCl, 3 mM MgCl_2_, 1 mM dNTPs, 5 mM DTT, and 10 U SUPERase·In (Invitrogen) for 1.5 h at 60 °C. The RNA was later hydrolyzed by 250 mM NaOH and the cDNA was circularized using CircLigase II (Lucigen). The circularized cDNA was shipped to the Scripps La Jolla Genomics Core for final library preparation with Illumina sequencing adaptors and sequenced using single-end sequencing on the Illumina NextSeq 500 platform.

### DMS MaPseq RNA Data Processing

DMS-MaPseq data processing followed a GitHub pipeline (https://github.com/borisz264/mod_seq). The adaptor sequence (GATCGGAAGAGCACACGTCTGAACTCCAGTCA) was trimmed from raw sequence via Skewer [45]. The first 5 nt, last 5 nt, and low quality nt were trimmed by Shapemapper 2.0. Reads were aligned to Saccharomyces cerevisiae 20S rRNA (Saccharomyces Genome Database), and read coverage and mutations at each position were counted by Shapemapper 2.0 to obtain the mutational rate at each nucleotide. The DMS accessibility for each individual A or C nucleotide was calculated by dividing the mutational rate for each A and C by the average mutational rate for all untreated A or C nucleotides, respectively. Thereafter, the values from the untreated samples were subtracted from the values for the DMS-treated samples, to obtained normalized DMS accessibility values for each nucleotide. Finally, the normalized DMS accessibility values from the two biological replicates were averaged.

Each residue was then sorted into one of four bins according to its normalized DMS accessibility score: 0.0-0.5, 0.5-1.0, 1.0-2.0, 2.0-4.0, and above 4. Residues that changed by one or more bins were considered to have different DMS accessibility.

Residues that are changed by one bin are indicated in yellow, residues changing one bin in one sample and two bins in one sample are shown in orange, and residues changing two bins in two samples are indicated in red.

### Antibodies

Antibodies against AFs were raised by Josman LLC in rabbits against purified recombinant protein, tested against recombinant protein and yeast cell lysate. Antibodies against Rps10 and Rps26 were raised in rabbits against chemically synthesized peptides by New England Peptide and tested against yeast cell lysate and recombinant protein (Rps26 only). The HA-antibody was from **Sigma Aldrich**.

## Acknowledgements

This work was supported by National Institute of Health grants R35-GM136323 and HHMI Faculty Scholar Grant 55108536 (to K.K.). We thank John McGrath (King’s College London) for providing us with information of the L216S mutation, and members of the Karbstein lab for discussion and comments on the manuscript. The authors declare no competing financial interests.

## Footnotes

1 Note that nonetheless yGFP-tagged Rps31 is found in the nuclei if 40S assembly is delayed [40]. Whether this reflects a weak association not captured by cryo-EM (*e.g.* by protein-protein interactions with Rps12), or is an artifact from the GFP fusion, or the stalled assembly is not clear.

## Supplementary Figures and Legends

**Figure S1:**
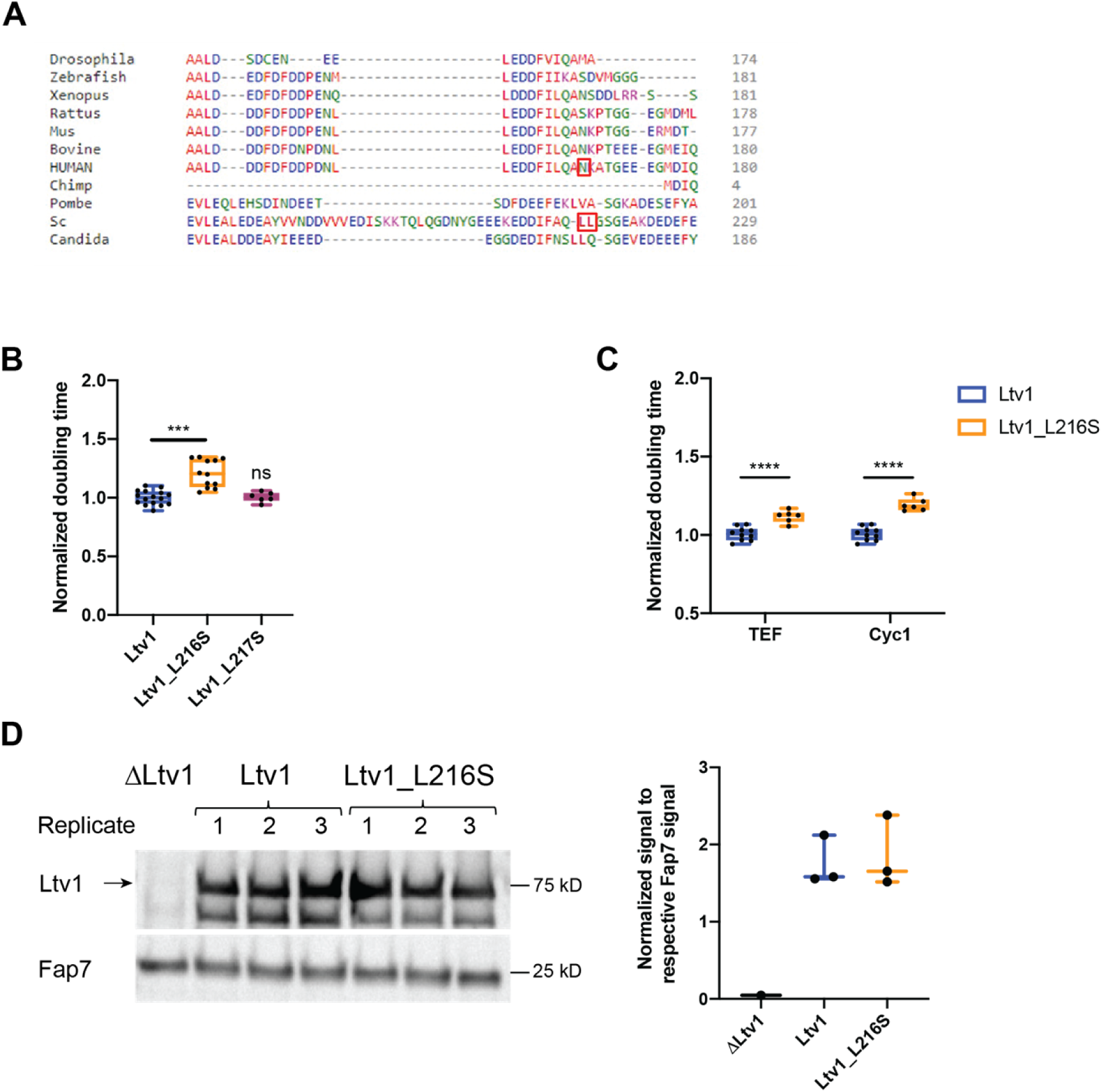
Yeast Ltv1_L216S corresponds to human Ltv1_N168S. (Relates to Figure 2) (A) Sequence alignment of a portion of Ltv1 highlighting N168 and L216 in human and yeast Ltv1, respectively. (B) Normalized doubling time of yeast lacking endogenous Ltv1 and expressing either wt Ltv1, Ltv1_L216S or Ltv1_L217S from TEF-promoter-driven plasmids. Significance was tested using an unpaired t-test. ***, P<0.001. Note that the data for WT Ltv1 and Ltv1_L216S are the same as in Figure 1B. (C) Normalized doubling time of yeast lacking endogenous Ltv1 and expressing either wt Ltv1, Ltv1_L216S or Ltv1_L217S from either TEF or Cyc1-promoter-driven plasmids. Significance was tested using an unpaired t-test. ****, P<0.0001. (D) Left: Western analysis of yeast lysates prepared from cells lacking endogenous Ltv1 and expressing either wt Ltv1 or Ltv1_L216S from TEF-promoter-driven plasmids. Three replicates for each are shown, as well as a control from ΔLtv1 cells. Fap7 is used as a loading control. Right: Quantification of the data on the left.

**Figure S2:**
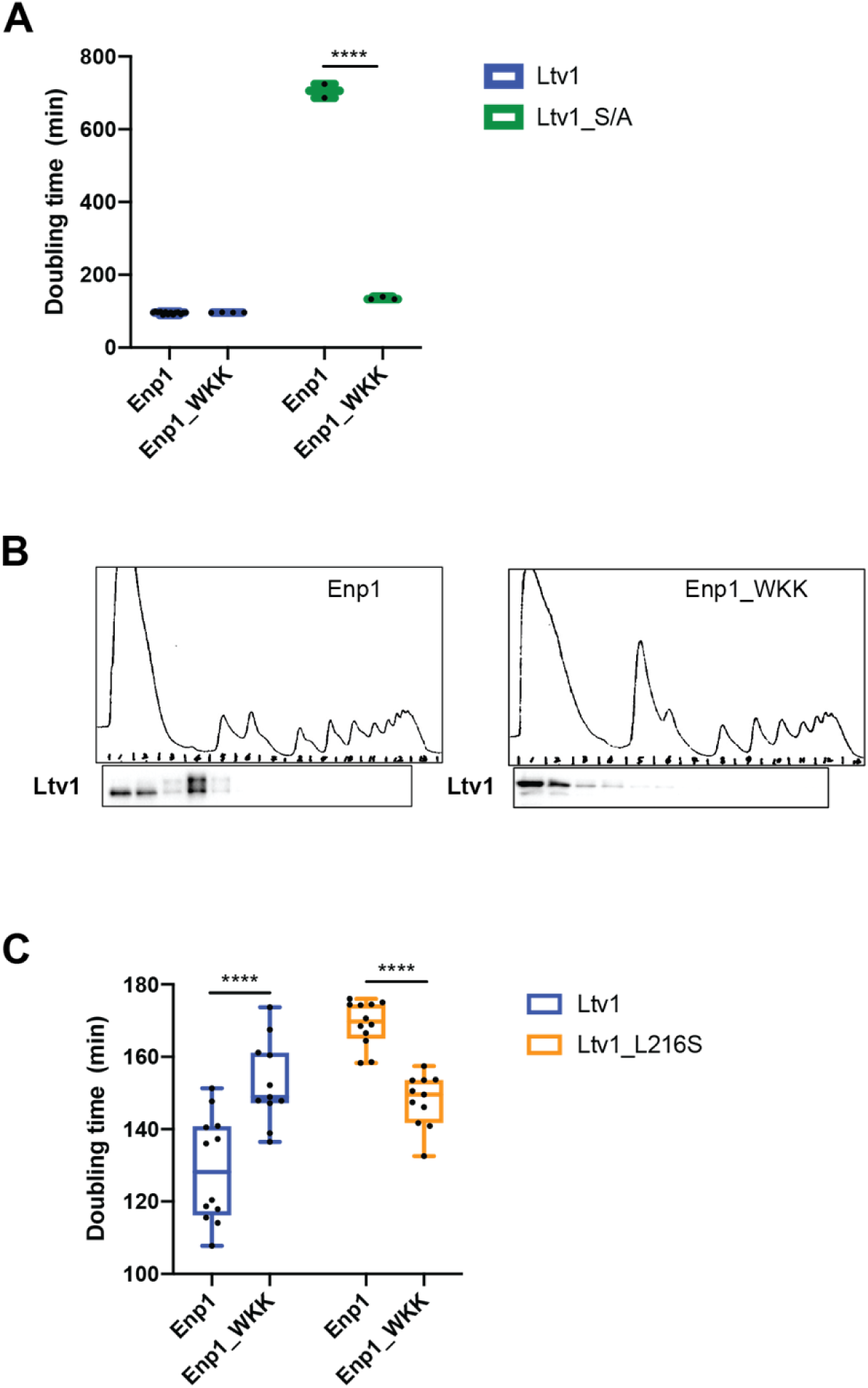
Enp1_WKK binds Ltv1 weakly, allowing for its phosphorylation independent release. (Relates to Figure 2) (A) Doubling time of yeast depleted for endogenous Enp1 and lacking endogenous Ltv1 and expressing either wt Enp1 or Enp1_WKK and wt Ltv1 or phosphorylation-deficient Ltv1_S/A from TEF-promoter-driven plasmids. Significance was tested using an unpaired t-test. ****, P<0.0001. (B) Absorbance profile (top) and Western blot (bottom) of yeast lysates from cells expressing either wt Enp1 or Enp1_WKK. (C) Doubling time of yeast depleted for endogenous Enp1 and lacking endogenous Ltv1 and expressing either wt Enp1 or Enp1_WKK and wt Ltv1 or Ltv1_L216S from TEF-promoter-driven plasmids. Significance was tested using an unpaired t-test. ****, P<0.0001.

**Figure S3:**
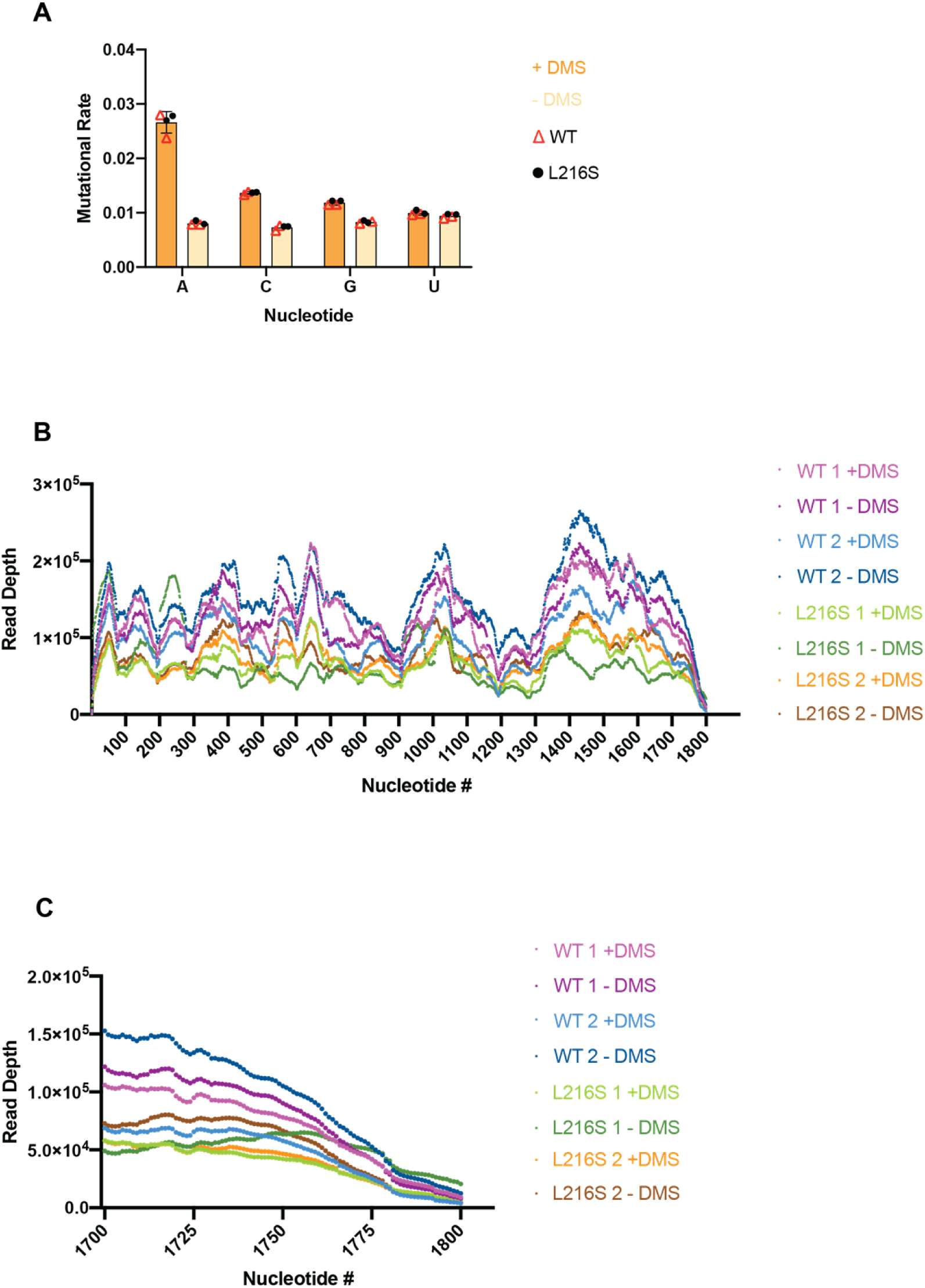
DMS-MaPseq quality control. (Relates to Figure 6). (A) Mutational rate changes for A and C residues upon DMS addition. Note that G also does get modified by DMS. (B) High read depth over the entire molecule. (C) Mature 18S rRNA is captured in all samples.

**Figure S4:**
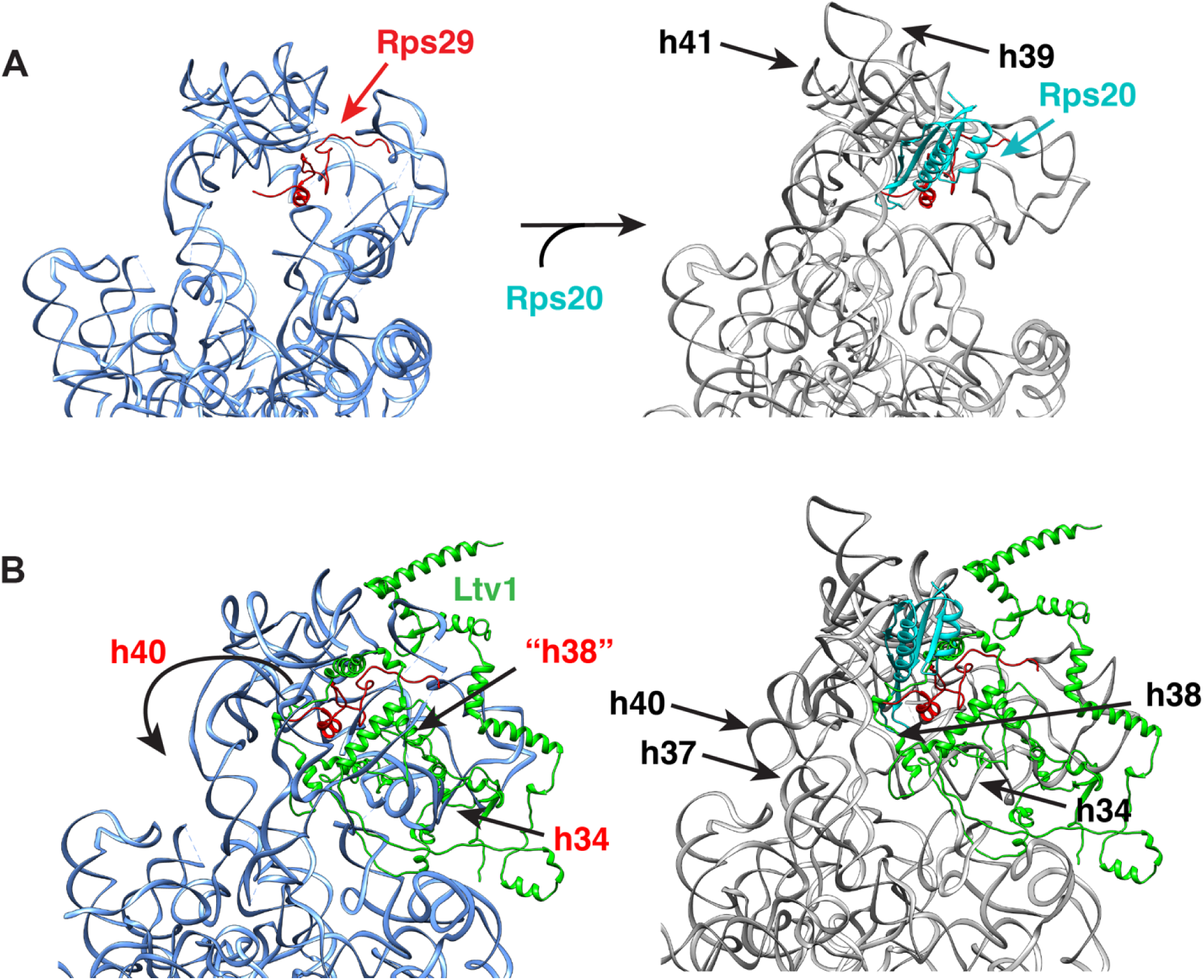
Binding of Rps20 is coupled to folding of h39 (Relates to Figure 6). (A) Structural detail of the 40S head (from PDB ID 7WTT) before and after Rps20 binding (from PDB ID 6FAI). For clarity only Rps20 and Rps29 are shown. (B) Structural details of the two 40S assembly intermediates in panel A, illustrating the sequential formation of helicase around j34-35-38 and the binding of Ltv1 adjacent to one strand of h38.

**Figure S5:**
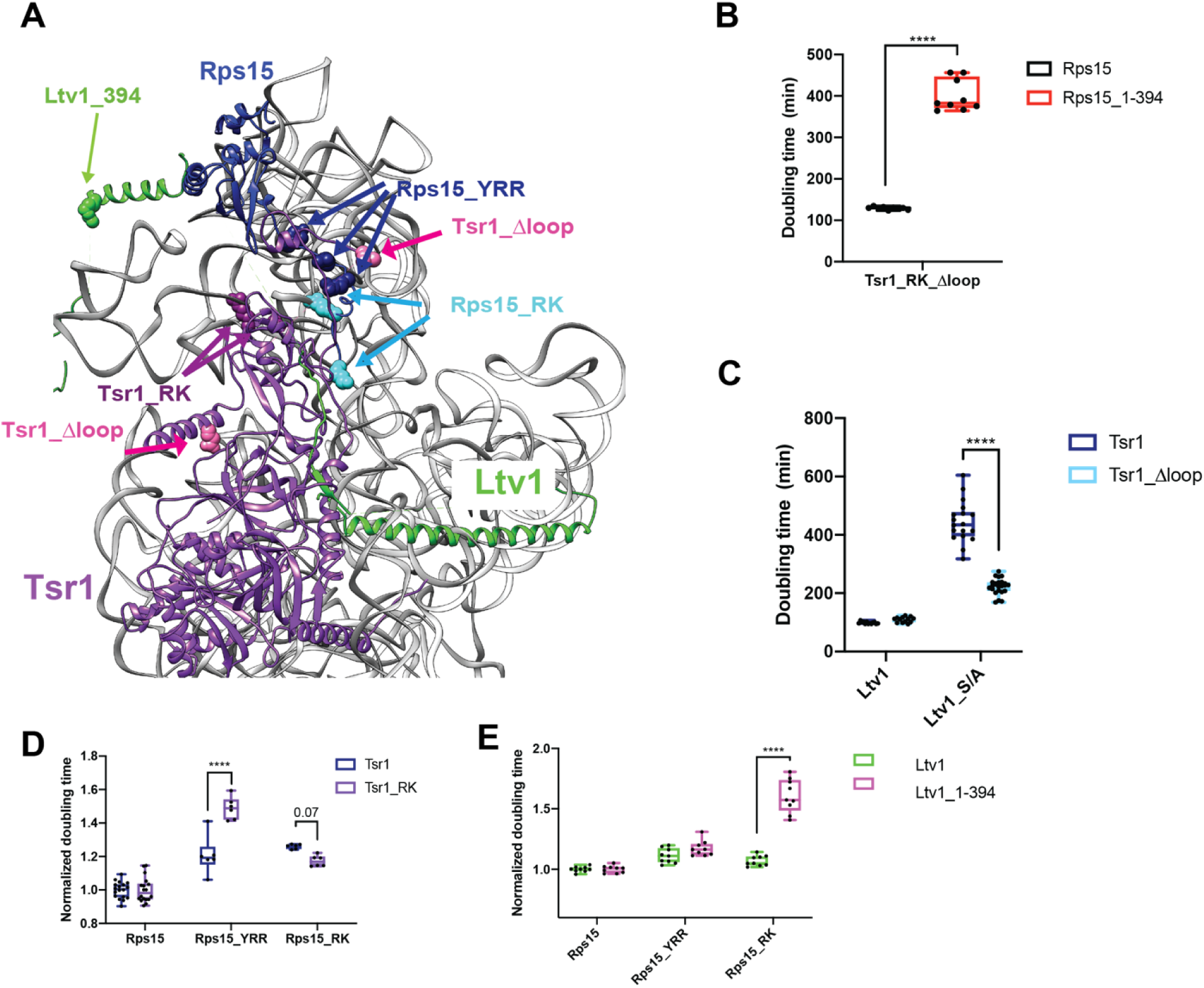
A structural and genetic network between Ltv1, Rps15 and Tsr1 (Relates to. Figure 7). (A) Structural detail of a composite structure of yeast pre-40S (PDB:6FAI) and human Ltv1 (PDB ID 6G18) highlighting elements in Tsr1, Rps15 and Ltv1. The residues mutated in Rps15_YRR and Rps15_RK are shown in blue and cyan spheres, respectively. The residues mutated in Tsr1_RK are shown in magenta spheres, and the first and last amino acids of the loop removed in Tsr1_Δloop are shown in magenta. The last amino acid in the Ltv1 truncation Ltv1_1-394 is shown in green space fill. (B) Doubling times for yeast cells expressing either wt Ltv1 or Ltv1_3-194, and Tsr1_RK_Δloop. Data for WT Ltv1 and Tsr1_RK_Δloop are in [21]. Significance was tested using an unpaired t-test. ****, P<0.0001 (C) Doubling times for yeast cells expressing either wt Ltv1 or the phosphorylation-deficient Ltv1_S/A and either wt Tsr1 or Tsr1_Δloop. Significance was tested using an unpaired t-test. ****, P<0.0001. (D) Doubling times (normalized to wt Rps15) for yeast cells expressing either wt Rps15, Rps15_YRR or Rps15_RK and wt Tsr1 or Tsr1_RK. Significance was tested using an unpaired t-test. ****, P<0.0001. (E) Doubling times (normalized to wt Rps15) for yeast cells expressing either wt Rps15, Rps15_YRR or Rps15_RK and wt Ltv1 or Ltv1_1-394. Significance was tested using an unpaired t-test. ****, P<0.0001.

**Table S1:** Residues with altered DMS accessibility.

| Residue No. | Average<br>WT Ltv1 | Average<br>Ltv1_L216S | $\Delta$(Ltv1_L216S<br>vs. WT) | Change |
| --- | --- | --- | --- | --- |
| 100 (A) | 0.9 | 1.6 | 0.7 | 2 bins, 1 bin |
| 103 (A) | 0.5 | 1.5 | 1.0 | 2 bins |
| 172 (C) | 0.5 | 1.8 | 1.2 | 1 bin |
| 184 (C) | 0.3 | 2.2 | 1.9 | 2 bins |
| 191 (C) | 5.1 | 2.7 | -2.4 | 2 bins, 1 bin |
| 221 (A) | 0.5 | 2.7 | 2.2 | 2 bins |
| 437 (A) | 0.5 | 1.7 | 1.2 | 2 bins |
| 990 (C) | 2.4 | 4.1 | 1.7 | 1 bin, unchanged |
| 1189 (A) | 0.4 | 1.4 | 0.9 | 2 bins |
| 1196 (A) | 0.6 | 1.4 | 0.8 | 1 bin, unchanged |
| 1197 (C) | 0.8 | 2.0 | 1.2 | 2 bins |
| 1505 (A) | 3.4 | 4.2 | 0.8 | 1 bin, unchanged |
| 1515 (A) | 1.2 | 2.2 | 1.0 | 1 bin, unchanged |
| 1591 (C) | 0.6 | 2.4 | 1.8 | 2 bins, 1 bin |
| 1753 (A) | 1.6 | 3.3 | 1.7 | 2 bins, 1 bin |

**Table S2:** **Yeast strains used in this study.**

| Strains | Description | Genotype | Reference |
| --- | --- | --- | --- |
| YKK73 | ΔLtv1 | BY4741 (MATa His3-1 Leu2-0 Met15-0 Ura3-0), Ltv1::KAN | [25] |
| YKK422 | ΔLtv1, Gal:Fap7 | BY4741, Ltv1::KAN, Gal:Fap7(NAT) | [46] |
| YKK1230 | ΔLtv1, Gal:S12 | BY4741, Ltv1::KAN; Gal:Rps12 (NAT) | This work |
| YKK1389 | ΔLtv1, snR35 | BY4741, Ltv1::NAT; snR35 (KAN) | [22] |
| YKK1576 | NOY504, ΔLtv1 | W303a (*MATα, leu2-3, 112, ura3-1, trp-1, his3-11, CAN1-100*); *rpa12::LEU2, Ltv1::HYG* | This work |
| YKK1115 | ΔLtv1, Gal:S29 | BY4741, Ltv1::HYG; Rps29B::KAN; Gal:3HA-Rps29 (NAT) | This work |
| YKK729 | ΔLtv1, Gal:S20 | BY4741, Ltv1::KAN, Gal:Rps20 (HYG) | [47] |
| YKK762 | ΔLtv1, Gal:S3 | BY4741, Ltv1::KAN, Gal:Rps3 (NAT) | [47] |
| YKK1117 | ΔLtv1, Gal:S15 | BY4741, Ltv1::KAN, Gal:Rps15 (NAT) | This work |
| YKK1625 | ΔLtv1, Gal:S31 | BY4741, Ltv1::NAT, Gal:Rps31 (HYG) | This work |
| YKK1190 | ΔLtv1, Gal:Rio2 | BY4741, Gal:Rio2(KAN), Ltv1::HYG | This work |
| YKK1142 | ΔLtv1, Gal:Enp1 | BY4741, Ltv1::KAN, Gal:Enp1(NAT) | [47] |
| YKK642 | ΔLtv1, Gal:Tsr1 | BY4741, Ltv1::KAN, Gal:Tsr1(NAT) | This work |
| YKK1184 | Gal:Rio2, Gal:Tsr1 | BY4741, Gal:Rio2(KAN), Gal:Tsr1(NAT) | [21] |
| YKK1444 | Gal:Rps31, Gal:Tsr1 | BY4741, Gal:31(HYG), Gal:Tsr1(KAN) | This work |

**Table S3:** Plasmids used in this study

| Plasmid | Description | Vector | Reference | Detailed description |
| --- | --- | --- | --- | --- |
| PKK3350 | WT Rio2 | TEF 413 | [21] |  |
| PKK3795 | Rio2_K105E | TEF 413 | [21] | K105E; see also [48] |
| PKK30356 | Rio2_loop | TEF 413 | [21] | R129A, H133A, R136A, R139A, D140A, K143A, K144A; see also [48] |
| PKK3295 | WT Tsr1 | TEF 416 | [31] |  |
| PKK3716 | Tsr1_RK | TEF 416 | [21] | R709E,K712E |
| PKK3895 | Tsr1_Δloop | TEF 416 | [21] | Substitution of amino acids 410 to 476 with PSSGSS; see also [49] |
| PKK30183 | WT Tsr1 | TEF 415 | [21] |  |
| PKK30184 | Tsr1_RK | TEF 415 | [21] | See PKK 3716 |
| PKK30116 | S15 | TEF 416 | [21] |  |
| PKK30180 | S15_YRR | TEF 416 | [21] | Y123I,R127K,R130K |
| PKK30181 | S15_RK | TEF 416 | [21] | R137E,K142E |
| PKK3890 | WT S20 | TEF 415 | [47] |  |
| PKK3934 | S20_DE | TEF 415 | This work | D113A,E115A |
| PKK3891 | S20_EYER | TEF 415 | [47] | E80K,Y82A,E83K,R85E |
| PKK3848 | TET S20 | pCM189 | This work | Tet-off Rps20 |
| PKK3606 | WT Ltv1 | TEF 413 | [46] |  |
| PKK3607 | Ltv1_S/D | TEF 413 |  | Ltv1_S336D,S339D,S342D |
| PKK30749 | Ltv1_L216S | TEF 413 | This work | Ltv1_L216S |
| PKK30753 | Ltv1_L217S | TEF 413 | This work | Ltv1_L217S |
| PKK30643 | Ltv1 | CYC 415 | This work |  |
| PKK30762 | Ltv1_L216S | CYC 415 | This work |  |
| PKK3693 | Ltv1_ΔC394 | TEF 413 | [21] | Deletion of residues after 394. |
| PKK30750 | Ltv1_L216S_S/D | TEF 413 | This work | Ltv1_L216S and Ltv1_S336D,S339D,S342D |
| PKK30752 | Ltv1_L216S_ΔC394 | TEF 413 | This work | Ltv1_L216S and deletion of residues after 394. |
| PKK30266 | Ltv1_YDY | TEF 413 | This work | Ltv1_Y82A,D83R,Y84A |
| PKK31167 | S31 | GPD 416 | This work |  |
| PKK31168 | S31_Δubi | GPD 416 | This work | Deletion of the N-terminal ubiquitin |
| PKK31169 | S31_ΔN | GPD 416 | This work | Deletion of the N-terminal extension, has ubiquitin |
| PKK3521 | WT S3 | TEF 416 | [47] |  |
| PKK3867 | S3_KK | TEF 416 | This work | S3_K7A,K10A |
| PKK3705 | S3_KR | TEF 416 | This work | S3_K75E, R76E |
| PKK4015 | TET Rps3 | pCM189 | [50] | Tet-off Rps3 |
| PKK30407 | WT S12 | TEF 415 | This work |  |
| PKK31165 | S12_R108 | TEF 415 | This work |  |
| PKK3944 | WT S29 | TEF 416 | This work |  |
| PKK3955 | TET S29 | pCM189 | This work | Tet-off Rps29 |
| PKK3541 | WT Enp1 | TEF 416 | [21] |  |
| PKK30159 | Enp1_WKK | TEF 416 | This work | Enp1_W224V,K228E,K231E |
| PKK3234 | WT 18S | pJV12 | [51] |  |
| PKK30582 | 18S_A1285A | pJV12 | [21] |  |
| PKK31082 | 18S_A1286A | pJV12 | This work |  |
| PKK30586 | 18S_A1288U | pJV12 | This work |  |
| PKK30974 | 18S_A1422U | pJV12 | This work |  |
| PKK30585 | 18S_G1288C | pJV12 | [21] |  |
| PKK30640 | 18S_C1327G | pJV12 | [21] |  |
| PKK30641 | 18S_G1288C+C<br>1327G | pJV12 | [21] |  |

